# Experimental and In Silico Analysis of the Structural Dynamics of Dengue NS2B-NS3 Protease

**DOI:** 10.64898/2026.09.02.748787

**Authors:** Susmida Seni Balakrishnan, Sivaraman Manjini, S Dhina, Kshirabdhi Dhal, Shibangi Das, Suresh Kumar Muthuvel

## Abstract

The Dengue virus (DENV), a major global health concern, causes dengue fever, predominantly affecting tropical and subtropical regions. The NS2B-NS3 protease complex of DENV is critical for viral replication, making it a promising target for antiviral drug development. This study investigates the structural dynamics of the NS2B-NS3 protease under varying pH conditions, integrating experimental and computational approaches. The recombinant NS2B-NS3 protease was expressed in E. coli, purified using affinity chromatography, and analysed for purity via SDS-PAGE. Circular dichroism (CD) spectroscopy revealed pH-dependent secondary structural changes, indicating stability at neutral to slightly basic pH and destabilization under acidic and highly basic conditions. Dynamic light scattering (DLS) analysis demonstrated protein aggregation and structural heterogeneity under extreme pH levels. Complementary in silico techniques, including homology modelling and molecular dynamics (MD) simulations, provided detailed insights into the conformational changes of the protease. The modelled structure, validated and refined through computational tools, revealed structural stability at physiological pH, with notable disruptions at pH extremes. DSSP (Dictionary of Secondary Structure of Proteins) and principal component analyses highlighted significant secondary structural transitions, especially at acidic pH where α-helices and β-sheets transformed into random coils Docking and MD simulation was carried out with the small molecule for therapeutic analysis between Dengue NS2bNS3 and small molecule. This study emphasizes the pH-dependent conformational plasticity of the NS2B-NS3 protease, contributing to the understanding of its functional mechanisms and providing a foundation for the rational design of pH-specific inhibitors. These findings underscore the importance of structural biology in advancing therapeutic strategies against dengue fever.

## Introduction

The DENV virus which causes Dengue fever, represents one of the numerous public health concerns and is prevalent mainly in tropical and sub-tropical regions. As estimated by the World Health Organization (WHO), There are up to 100 - 400 million cases of Dengue fever and around 50 percent of the population is under threat [1]. Four serotypes of Dengue viruses exist namely: DENV-1, DENV-2, DENV-3, DENV-4. Infection by any one of these serotypes could cause symptoms of mild fever to severe dengue hemorrhagic fever and Dengue shock syndrome, especially in secondary infections of young children who inherited maternal antibodies or immuno-compromised adults [2]. Even with dedicated research, antiviral therapies against all 4 serotypes are not currently available hence, raising the need for identifying new therapeutic targets and antiviral drugs for treatment against DENV infections are of high priority. The DENV NS2B-NS3 protease complex is excellent target for development of antiviral therapy against this virus. This complex plays a significant role in the processing of viral polyprotein precursor into mature proteins prior to viral replication [3]. The full-length NS3 protein (∼618 amino acids; aa) has multiple roles; it is a protease within the N-terminal ∼185 aa and an RNA helicase, as well as a nucleoside triphosphatase within the rest of NS3 protein [4].The protease functions to cleave the viral polyprotein at specific sites to release mature NS3 protease complex. The region from ∼172 to yje C-terminus harbors ATP-dependent RNA helicase and the 5’ RNA triphosphatase activities that are involved in viral RNA replication and 5’-caping [6]. Although for viral replication and 5’ cap addition, full-length NS2B and NS3 proteins are required, for in vitro protease activity, conserved ∼48aa hydrophilic domain of NS2B and ∼ 185aa of the N-terminal region of NS3 protease forming an active serine protease complex are sufficient [8]. NS2B cofactor in structural investigations appeared to interact with the NS3 protease domain, where it formed part of a β-hairpin structure that increases enzyme catalytic efficiency [5]. The objective of this work is to investigate the structural changes of NS2B-NS3 during variations in pH playing a role during the life cycle of the dengue virus. To appreciate how viruses multiply and cause diseases, it is necessary to understand that there are proteins that are stable, fold and carry out functional activities that can be gleaned from structural changes of proteins as a function of pH [7] .Understanding the structural dynamics of the NS2B-NS3 complex under different physiological conditions, such as variations in pH. Clarifying the conformational variations of viral proteins can reveal important information about their modes of action because these proteins must maintain stability and functional activity in a variety of intracellular-milieu. By investigating its stability and conformational shifts, it will contribute to a deeper understanding of its functional mechanism [9,10,12]. This study emphasizes the importance of structural biology and bioinformatics in solving dengue fever’s global challenge and moving forward with the development of innovative antiviral treatments.

## Materials and Methods

### Bacterial Strain and Expression Plasmid

The DENV2 were expressed from 2L cultures of TOP10F′ cells (Invitrogen) transformed with the expression plasmids pQE30-NS2BH(QR)-NS3pro [13]. The DENV1 and DENV4 NS2BH-NS3 expression plasmids were created by PCR amplification of coding sequences from the full-length cDNA clones of DENV1 and DENV4, each of which contained the credible C-terminal amino acids of NS2B tailed by the NS3pro domain. A clone of DENV3 cDNA in pRS424 vector (YU4N) with a deletion of E and a portion of NS1 genes that could be sustainably reproduced in E. coli (Gifted by Dr. Barry Falgout, FDA) was utilized for overlap extension PCR using standard molecular biology methods to construct the DENV3 NS2BH-NS3pro plasmid. The DENV1, DENV3, and DENV4 plasmid clones for the NS2BH-NS3pro domains included the four amino acids from the NS2B C-terminus linked to NS3pro, which includes the NS2B-NS3 cleavage site in pET-32a between the NcoI and BamHI restriction sites (unpublished data). By surrounding the two amino acid residues (QR), at the NS2B-NS3 location, the NS3pro domain and the NS2B hydrophilic domain are encoded by the DENV2 protease expression plasmid. NS2B-NS3pro expression plasmids and DENV1-4-transformed E. coli cells were cultured in LB medium supplemented with ampicillin (100 μg/mL) and 0.1% glucose at 37°C until the OD600 reached 0.6 – 0.7.

### Bacterial Culture and Protein Expression

NS2B-NS3pro expression plasmids and DENV1-4-transformed E. coli cells were cultured in LB medium supplemented with ampicillin (100 μg/mL) at 37°C. The culture’s optical density was monitored at 600 nm (OD600), and when the OD600 reached 0.4 – 0.5, protein expression was induced by adding isopropyl-β-D-thiogalactopyranoside (IPTG) to a final concentration of 0.5 mM. The cultures were incubated for an additional 3 – 4 hours at 37°C with shaking. After induction, bacterial cells were harvested by centrifugation at 5000 rpm for 15 minutes at 4°C. The resulting cell pellet was stored at -80°C until further use.

### Protein Purification

The cell pellet was resuspended in Buffer A (50 mM Tris-HCl, pH 7.5, 150 mM NaCl) and lysed by sonication for 10 minutes at a frequency of 40 Hz. The lysate was clarified by centrifugation at 8000×g for 1 hour at 4°C. The supernatant was then subjected to nickel affinity chromatography using a pre-equilibrated Ni-NTA resin (3 ml suspension) with Buffer A. Proteins were eluted stepwise with Buffer A containing increasing concentrations of imidazole (50 mM to 250 mM).

### SDS-PAGE Analysis

The elution fractions were analysed by SDS-polyacrylamide gel electrophoresis (SDS-PAGE) to confirm the presence and purity of the target protein. The elution profile was monitored for effective recovery of the recombinant protease.

### Circular Dichroism (CD) Spectroscopy

The auxiliary structure of the dengue NS2B-NS3 protease was analysed utilizing circular dichroism (CD) spectroscopy. The protein at various pH levels were tested test with the Tris buffer. The protein concentration was balanced to around 0.8 mg/ml, measured by UV absorbance at 280 nm. CD spectra were recorded in a far-UV extend to 190 – 260 nm utilizing a Jasco J-815 spectropolarimeter at room temperature. A quartz cuvette with a way length of 1 mm was utilized. Each measurement was an average of three scans, with standard correction performed using the corresponding buffer. The data was expressed as mean residue ellipticity (MRE) in deg·cm² dmol⁻¹

### Dynamic Light Scattering (DLS) Analysis

Dynamic Light Scattering (DLS) was performed to evaluate the hydrodynamic measure dissemination and accumulation state of the dengue NS2B-NS3 protease. Estimations were conducted utilizing a Malvern Zetasizer Nano S at 25°C. Protein samples were prepared at a concentration of 0.8 mg/ml in 50 mM Tris-HCl (pH 7.5) and 150 mM NaCl. Prior to analysis, the samples were filtered through a 0.22 µm syringe filter to remove particulates. DLS measurements were performed in a disposable microcuvette at a scattering angle of 173°. The Z-average hydrodynamic diameter (d.nm), polydispersity index (PdI), and size distribution by intensity were recorded.

### Dataset Preparation for *in silico* approach

The three-dimensional structure of the Dengue infection NS2B-NS3 protease complex was modelled using I-Tasser and the structure refinement was done using modLoop. The structure validation was done using Ramachandran Plot (*Supplementary data S1*). Visualization and auxiliary investigation of the protein were performed using PyMOL and UCSF ChimeraX. These visualization tools were utilized to create two-dimensional and three-dimensional representations of the protein complex for auxiliary examination [14,15].

### Protonation States Examination (H++ Server)

Protonation states of the protein’s ionizable buildups were decided utilizing the H++ web server, which predicts the pK values of acidic and fundamental bunches based on the natural pH. The H++ server too completes the structure by including lost hydrogen molecules and yields the comes about in broadly acknowledged groups, counting PDB and PQR. These protonation states are pivotal for understanding macromolecular structure-function connections affected by electrostatics [16].

### Molecular Docking and Molecular Dynamic (MD) Simulations

The Molecular docking studies were carried out using the Schrödinger Maestro suite to study about the binding interactions of isatin with the Dengue NS2B–NS3 protease under different pH conditions (acidic to basic). The protein and ligand structures were prepared and docking analysis was performed using the standard Schrödinger docking workflow to predict the binding affinity and interaction pattern of the ligand. The docked complexes were further analysed using LigPlot+ to generate 2d interaction images.Molecular Dynamic (MD) reenactments were conducted to explore the conformational elements and auxiliary solidness of the protein[45]. The reenactments were performed utilizing the Groningen Machine for Chemical Recreations (GROMACS 2021) bundle [17,18]. The protein topology was produced with the pdb2gmx module, utilizing the GROMOS54a7 drive field [19]. Vitality minimization was accomplished utilizing the steepest plummet calculation with a drive merging model (Fmax) of 1000 kJ/mol·nm over 5000 steps. The recreation framework was neutralized with NaCl particles and solvated utilizing the SPC/E water show in a triclinic box. Equilibration was carried out in two stages: NVT (consistent number of particles, volume, and temperature) and NPT (steady number of particles, weight, and temperature), each for 5000 steps. The temperature was kept up at 300 K, and weight at 1 bar, utilizing the Berendsen weight coupling strategy amid NPT equilibration. Taking after equilibration, a 100 ns generation recreation was performed utilizing the leap-frog integrator. Trajectory investigation was carried out utilizing GROMACS devices, assessing parameters such as root mean square deviation (RMSD), root mean square variance (RMSF), and hydrogen holding. To examine collective movements and overwhelming conformational changes in the protein amid the MD reenactment, Principal Component Analysis (PCA) was performed utilizing the GROMACS gmx covar and gmx anaeig apparatuses. The nuclear facilitates of the Cα iotas from the recreation direction were utilized to build the covariance lattice. Eigenvalues and eigenvectors were calculated to distinguish the vital components speaking to the biggest conformational fluctuations. The to begin with three foremost components (PC1, PC2 and PC3) were plotted to visualize the overwhelming movements. The comparing eigenvector projections were analysed to decide the adaptability and portability of the protein amid the reenactment [20,21].

### Secondary Structure Examination (DSSP)

The GROMACS DSSP module was used to monitor changes in the secondary structure elements of the protein during simulations. This analysis categorized secondary structure patterns into α-helix, β-sheet, β-bridge, coil, turn, twist, and other conformations. Additionally, secondary structure variability was determined according to the standard DSSP algorithm [11].

## Result and Discussion

### Expression and Protein Purification

Dengue NS2B-NS3 clone was successfully expressed and purified the recombinant Dengue NS2B-NS3 protein using Escherichia coli and Nickel NTA affinity chromatography. The purification process involved a buffer A at pH 7.4. During the purification, we used an imidazole gradient to elute the protein from the Nickel column and collected various fractions for further analysis. To assess the purity of the collected fractions, we conducted sodium dodecyl sulfate-polyacrylamide gel electrophoresis (SDS-PAGE) [23]. The results showed a prominent band at around 29 kDa, which corresponds precisely to the expected molecular weight of the NS2B-NS3 fusion protein with BSA as a marker (***Fig.1***).

**Fig. 1.**
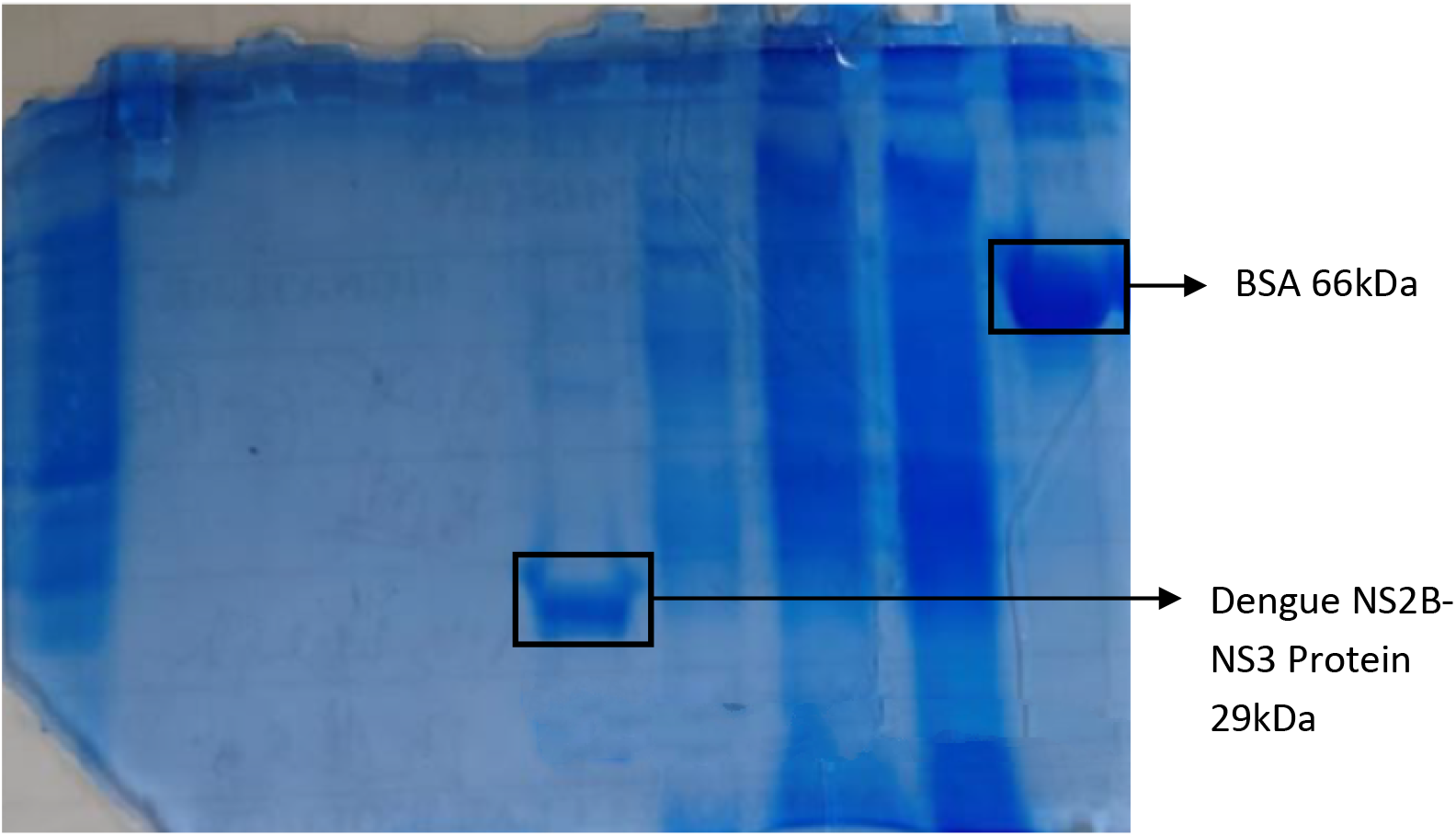
SDS-PAGE analysis of the purified Dengue NS2B-NS3 protein. Purified Dengue NS2B-NS3 protein was analysed by SDS-PAGE. Lane 1: BSA as marker (66kDa), Lane 2: After sonication Lane 3: Supernatant after centrifugation, Lane 4: Flow through. Lane 5:100mM. The major band at Lane 5 confirming successful expression and purification.

The molecular weight reflects the combined size of the NS2B cofactor and the NS3 protease, and the lack of additional bands indicates that our purification process was successful, yielding a protein that is mainly intact with minimum degradation or non-specific binding. Interestingly, we opted to skip the dialysis step, a common practice in protein purification aimed at removing salts and other small molecules that could interfere with future applications [24]. We carefully controlled the salt and imidazole concentrations to reduce any potential negative impact on downstream analyses, following successful examples in similar recombinant protein studies [25]. By maintaining a salt concentration, we ensured the protein remained stable. The SDS-PAGE results confirmed the absence of significant contaminants. The clear 29 kDa band indicates that the protein is not only pure but also properly folded and fully expressed.

### Circular Dichorism (CD) Spectroscopy

The PDI values varied across the pH range, reflecting the heterogeneity of the protein population. Acidic pH 4 - 6 were associated with higher PDI values (below 0.5), indicative of significant aggregation and conformational heterogeneity. Neutral pH 7.5 displayed a relatively stable distribution, with a moderate PDI value was 0.495. At basic pH, while pH 8 showed extremely high heterogeneity value of PDI was 0.508. At pH 9 show that Dengue NS2b-NS3 protein predominantly forms a stable population with a hydrodynamic diameter of 190.3 nm and a PDI of 0.338, indicating moderate heterogeneity. And pH 10 exhibited a moderate level of polydispersity PDI value was 0.721, suggesting partial stabilization under strongly alkaline conditions.

- ***Impact of pH on Protein Stability***

The DLS data highlights the influence of pH on the stability and aggregation behavior of the Dengue NS2B-NS3 protease:

- **Acidic pH (pH 4 - 6):** Increased protonation of acidic residues destabilizes intramolecular interactions, leading to partial unfolding and aggregation. This is consistent with studies demonstrating that low pH disrupts salt bridges and hydrogen bonding networks.

The circular dichroism (CD) spectra data under varying pH conditions provide valuable insights, how the secondary structure of the protein changes across both acidic and basic environments. CD spectroscopy helps monitor the ellipticity values, especially around the characteristic wavelengths for alpha-helices (around 208 nm) and beta-sheets (around 218 nm), which are indicative of the structural stability and folding of the protein. Measurements were taken across a wide range of pH levels from acidic (pH 4 - 6) to basic (pH 8 - 10), with neutral pH 7.5 allowing us to examine the influence of pH on protein conformation. The gradual trend in ellipticity from acidic to basic pH levels suggests that extreme pH conditions (both very low and very high) can destabilize the protein, while near-neutral to mildly basic conditions favor structural stability. At neutral to slightly basic (pH 7.5 and 8), the protein maintains a stable conformation, with contributions from both alpha-helices and beta-sheets, indicating an intact secondary structure. Under acidic conditions (pH 4 - 6), the observed structural instability may be attributed to protonation effects on acidic residues (***Fig.2***), which can interfere with stabilizing hydrogen bonds and ionic interactions. Such effects are consistent with studies that show acidic environments can disrupt proteins with high acidic residue content, leading to partial unfolding [26, 27].

**Fig. 2.**
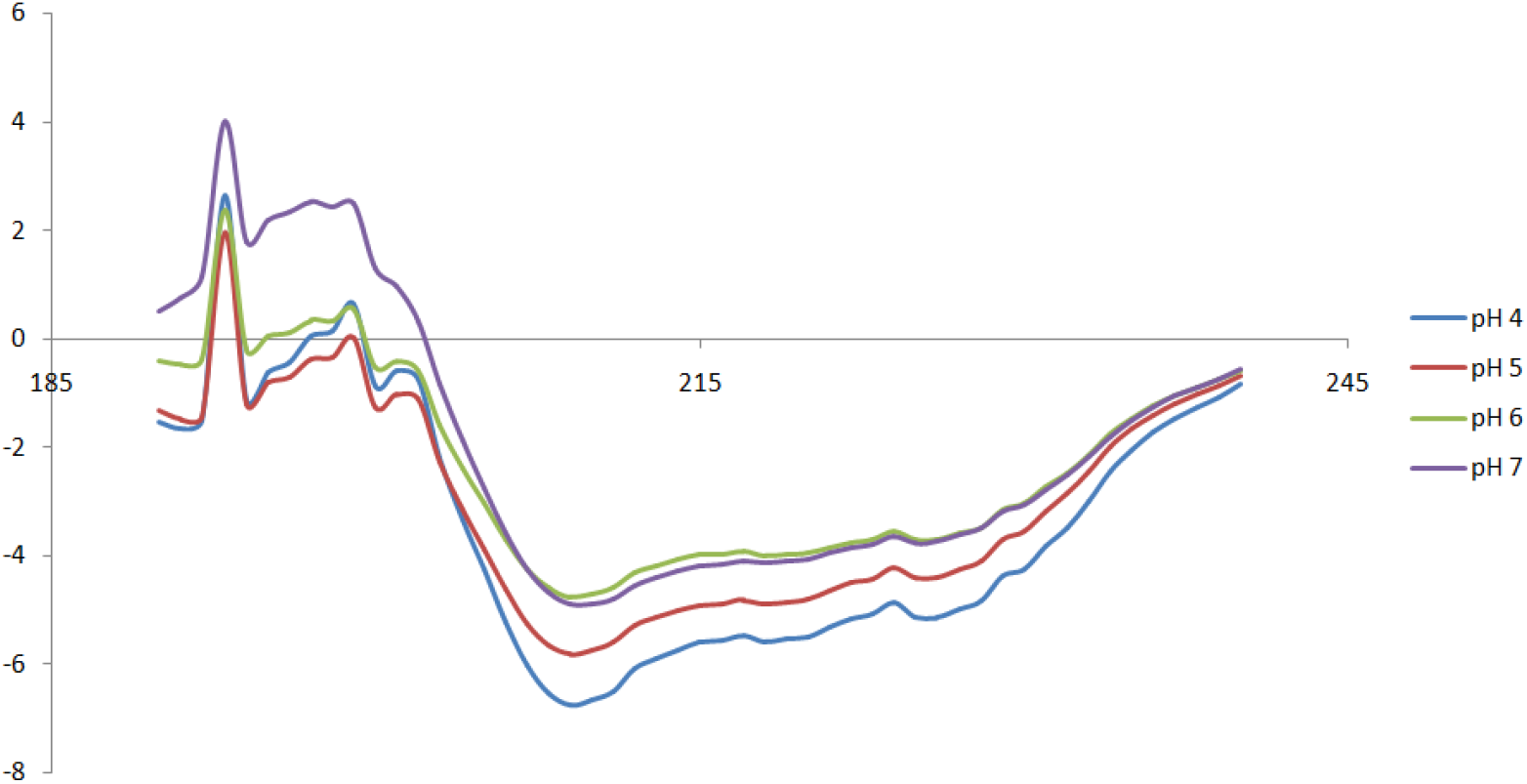
CD spectra of the Dengue NS2B-NS3 protein were recorded to assess secondary structure under acidic (pH 4 - 6) and neutral (pH 7.5) conditions. Spectra were measured in the far-UV region (190 – 260 nm) using a 0.1 cm path length cuvette.

In highly basic conditions (pH 8 - 10), while the protein remains mostly stable, certain ionizable groups on the protein may undergo deprotonation (***Fig.3***). This can occasionally lead to minor structural adjustments, as some hydrogen bonding and electrostatic interactions are altered. The stability of the protein’s secondary structure at near-neutral and slightly basic pH 7.5 and 8 suggests that these conditions may be ideal for maintaining its functional conformation. In contrast, extreme pH environments (both acidic and highly basic) lead to structural changes that could impact protein function, particularly if such conformational changes interfere with the protein’s active site or interaction surfaces.

**Fig. 3.**
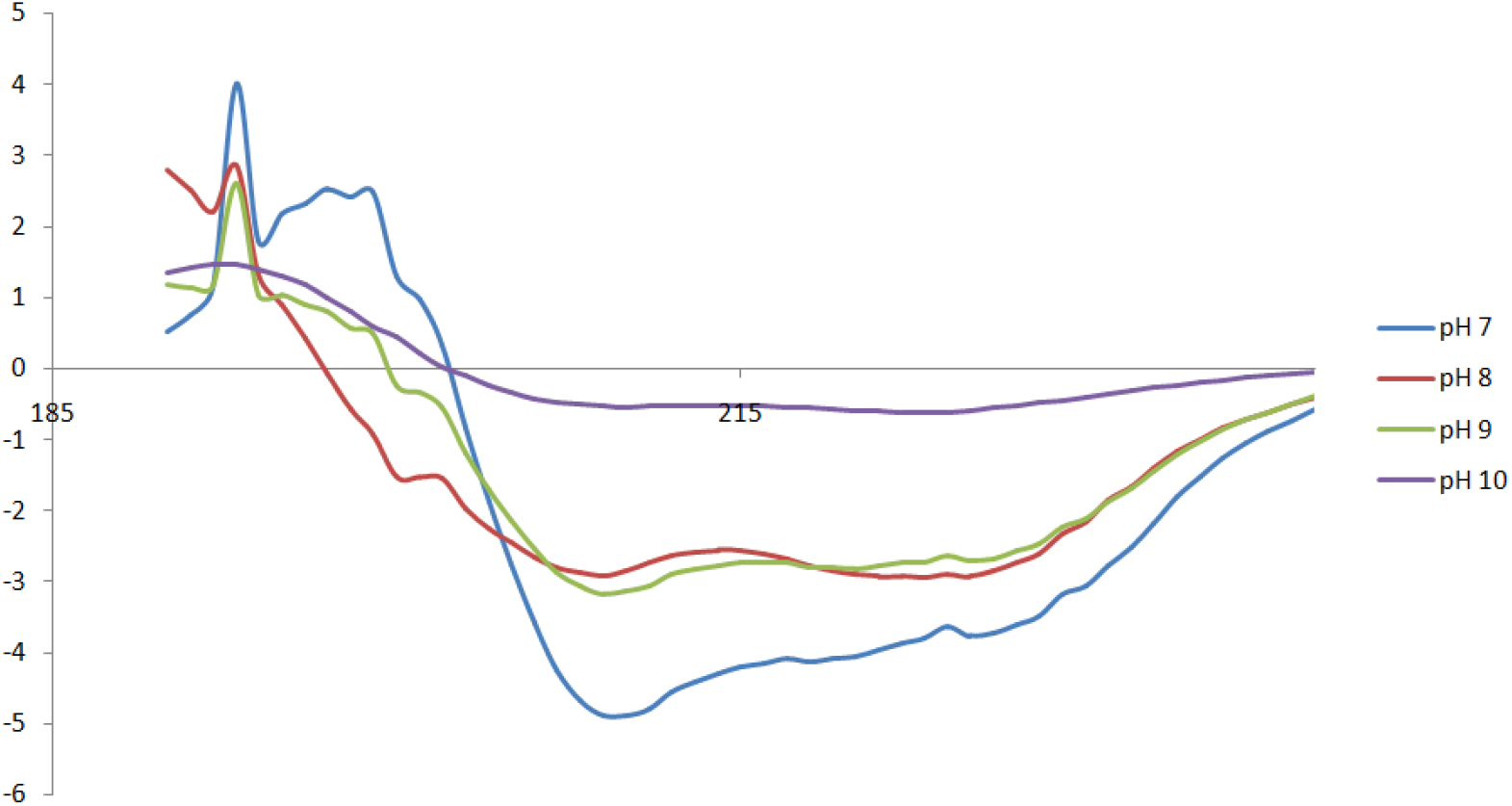
CD spectra of the Dengue NS2B-NS3 protein were recorded to assess secondary structure under acidic (pH 8 – 10) and neutral (pH 7.5) conditions. Spectra were measured in the far-UV region (190 – 260 nm) using a 0.1 cm path length cuvette.

The structural stability of the Dengue NS2B-NS3 protease was assessed using CD spectroscopy under varying pH conditions (4 - 10). The CD spectra and secondary structure analysis revealed distinct trends in protein folding and stability.

### Acidic pH (pH 4 – 6)

The CD spectrum showed significant deviations at pH 4 and 5, characterized by reduced helicity (27.6% at pH 4 and 25.1% at pH 5) and a complete absence of β-sheets (0% for both). Random coil content was notably high (37.9%), indicating partial unfolding. At pH 6, the secondary structure slightly stabilized with an increase in helicity (28.4%) but still exhibited destabilization compared to pH 7.5.

### Neutral pH (pH 7.5)

CD spectra at pH 7.5 displayed the highest structural stability, with 33.2% helicity and a balanced distribution of secondary structure components. This highlights that the protease retains its native conformation under physiological conditions.

### Basic pH (pH 8 – 10)

At pH 8, a slight destabilization occurred, with helicity decreasing to 29.2% and random coil content increasing to 47.3%. However, the protease retained significant structural integrity. The spectrum at pH 9 showed a helical content of 31% but still displayed flexibility with moderate random coil and turn structures. At pH 10, a drastic increase in helicity (82.1%) and a reduction in random coil (17.9%) were observed. Though, this apparent stabilization might indicate misfolding or structural rearrangement, as corroborated by the highest RMSD value (87.955), suggesting significant conformational deviation. These findings corroborate previous studies showing that environmental pH significantly influences the stability and activity of viral proteases ***(Table.S2)***. For instance, the NS3 protease from hepatitis C virus also exhibits optimal activity near physiological pH, with destabilization under extreme conditions [40]. This pH-dependent plasticity may have implications for the design of inhibitors targeting specific conformational states of the protease.

### Dynamic Light Scattering (DLS)

Dynamic Light Scattering (DLS) analysis was conducted to assess the hydrodynamic size distribution, polydispersity, and aggregation state of the Dengue NS2B-NS3 protease under varying pH conditions from acidic to basic (pH 4 - 10). This investigation provides insights, how pH influences protein stability, conformation, and aggregation behavior. Key findings and interpretations are discussed.

- Hydrodynamic Radius and Size Distribution

At pH 4, the DLS data revealed single peaks, indicating a homogeneous size distribution. The presence of broad peaks suggested that the protein was prone to aggregation under acidic conditions. This was attributed to the disruption of stabilizing electrostatic interactions and hydrogen bonding due to protonation of acidic residues. The aggregation behavior observed aligns with reports that low pH induces partial unfolding and exposure of hydrophobic regions, promoting intermolecular interactions and aggregate formation [26,28]. The protein sample exhibited single distinct peaks in the intensity-weighted size distribution. The Z-average size was recorded at 234.4 nm with peaks at ∼234.4 nm, PDI of 0.050 [29]. The presence of these peaks suggests a mixture of monomeric species and larger aggregates. The emergence of larger aggregates indicates enhanced aggregation tendencies at this pH. At pH 5, the Z-average diameter, which represents the overall mean particle size in solution, was recorded as 101.3 nm, 454.6 nm, and 5172 nm. The polydispersity index (PDI) was measured at 0.447, with a Z-average diameter of 165.1 nm. At pH 6, the Z-average hydrodynamic diameter was 276.4 nm, with a prominent peak at 169.8 with the intensity percentage of ∼89.7. The broad size distribution indicated by the high polydispersity index PDI of 0.303, suggests that the protein undergoes partial aggregation under mildly acidic conditions. This aggregation can be attributed to the disruption of ionic interactions and partial unfolding of the protein, consistent with previous studies on protein stability in acidic environments [26]. At neutral pH, the sample displayed a predominant peak centered at ∼63.71 of intensity percentage, corresponding to the monomeric or minimally aggregated state of the protein. The Z-average diameter was 178.0 nm, with a PDI of 0.926, indicating moderate polydispersity. The primary peak accounted for the majority of the sample intensity, suggesting that the protein maintains relative structural stability and solubility in neutral conditions. This stability aligns with reports that neutral pH favors the maintenance of intramolecular hydrogen bonding and electrostatic interactions. At pH 8, the Z-average size increased significantly to 5255 nm, with a PDI of 0.508, indicating extensive aggregation and heterogeneity in the sample. The size distribution revealed three peaks, with the dominant peak at ∼174.9 accounting for 73.2% of the intensity. The high Z-average and PDI values suggest that alkaline conditions favor aggregation, likely due to changes in charge distribution and reduced solubility of the protein under these conditions. This finding aligns with studies indicating that pH near the protein’s isoelectric point can promote aggregation [30]. At pH 9 indicate that the Dengue NS2b-NS3 protein exists predominantly as a stable population with a hydrodynamic diameter of 190.3 nm (Peak 1). The observed PDI (0.338) and minor secondary peak suggest a small degree of heterogeneity, possibly due to the pH-induced structural or conformational changes. The absence of significant aggregation points to the structural stability of the protein at alkaline pH, consistent with its functional role under physiological conditions [22]. At pH 10, the Z-average diameter was 501.3 nm, with two major peaks at ∼212.7 nm and ∼38.57 nm. The PDI was 0.721, indicating moderate polydispersity. The relatively lower PDI compared to pH 8 suggests a reduction in aggregation and better particle dispersion at pH 10 (***Fig.4***). The negative charge at this pH may enhance electrostatic repulsion, mitigating large-scale aggregation. However, the presence of smaller peaks indicates that transient aggregation still occurs [31].

**Fig. 4.**
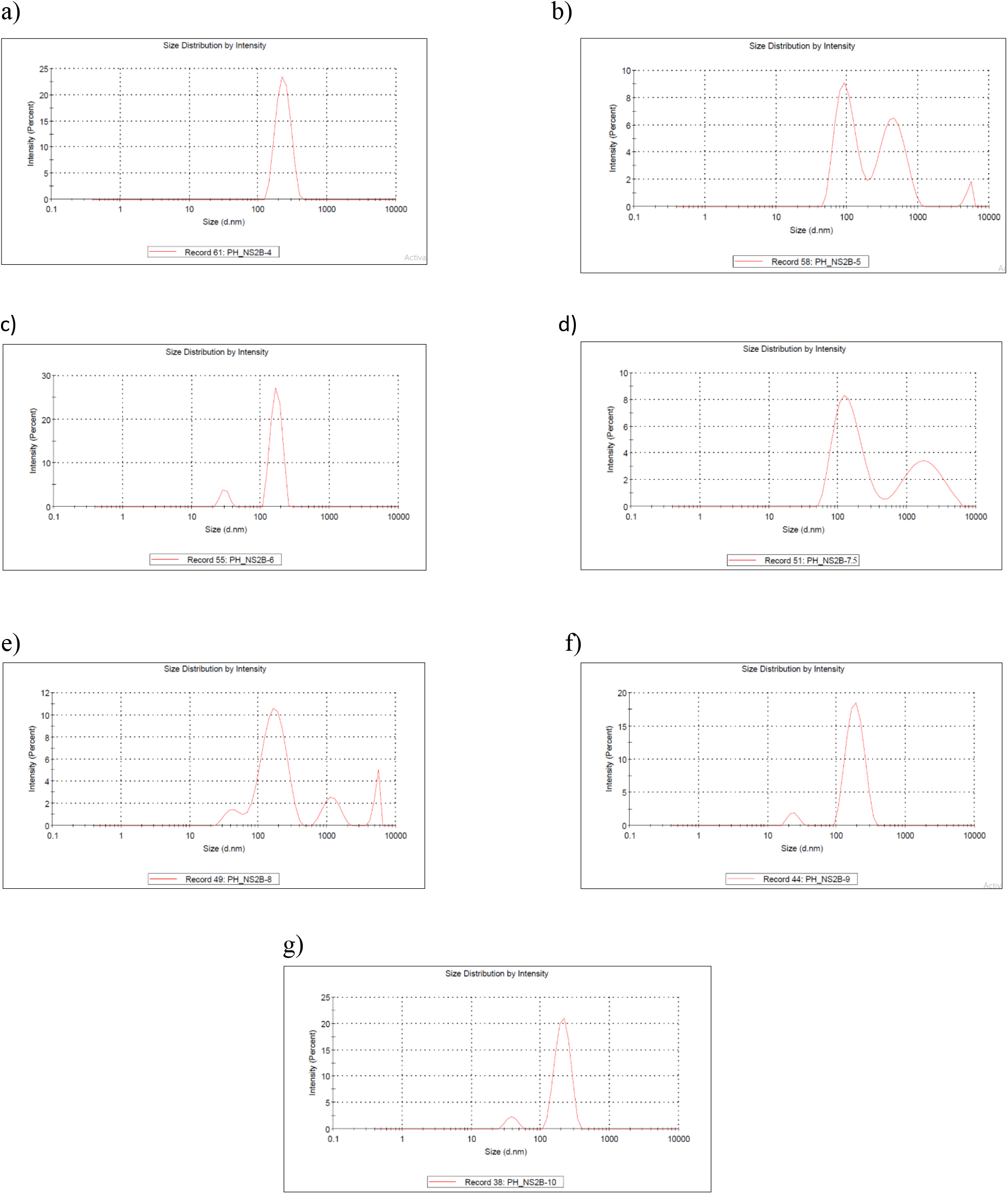
Dynamic Light Scattering (DLS) analysis of Dengue NS2B-NS3 protein under varying pH conditions: (a) pH 4, (b) pH 5, (c) pH 6, (d) pH 7.5 (e) pH 8 (f) pH 9 and (g) pH 10. The data illustrate the size distribution and aggregation behavior of the protein across acidic, neutral, and alkaline environments. Polydispersity Index (PDI) and Aggregation Behavior

The PDI values varied across the pH range, reflecting the heterogeneity of the protein population. Acidic pH 4 - 6 were associated with higher PDI values (below 0.5), indicative of significant aggregation and conformational heterogeneity. Neutral pH 7.5 displayed a relatively stable distribution, with a moderate PDI value was 0.495. At basic pH, while pH 8 showed extremely high heterogeneity value of PDI was 0.508. At pH 9 show that Dengue NS2b-NS3 protein predominantly forms a stable population with a hydrodynamic diameter of 190.3 nm and a PDI of 0.338, indicating moderate heterogeneity. And pH 10 exhibited a moderate level of polydispersity PDI value was 0.721, suggesting partial stabilization under strongly alkaline conditions.

- Impact of pH on Protein Stability

The DLS data highlights the influence of pH on the stability and aggregation behavior of the Dengue NS2B-NS3 protease:

- **Acidic pH (pH 4 - 6):** Increased protonation of acidic residues destabilizes intramolecular interactions, leading to partial unfolding and aggregation. This is consistent with studies demonstrating that low pH disrupts salt bridges and hydrogen bonding networks.
- **Neutral pH (pH 7.5):** The protein exhibited minimal aggregation, likely maintaining its native-like conformation, as neutral conditions support optimal electrostatic interactions and solubility.
- **Basic pH (pH 8 - 10):** Alkaline conditions resulted in varying levels of aggregation. While pH 8 favored extensive aggregation, the reduced aggregation at pH 10 suggests enhanced repulsion among negatively charged particles, partially stabilizing the sample [30,31].

### Sequence Alignment and Homology Modelling of Dengue NS2B-NS3 Protein

The structural model of the Dengue virus NS2B-NS3 protease was successfully generated to elucidate its molecular conformation and functional insights. The sequence of the NS2B-NS3 protein was obtained and aligned against homologous structures using BLASTp and Clustal Omega. Sequence alignment revealed significant conservation in the protease catalytic triad (His, Asp and Ser residues) and NS2B cofactor-binding region, essential for protease activity. This conservation provided a reliable foundation for homology modelling. The primary structure of NS2B-NS3 was submitted to the I-TASSER tool, which generated a preliminary three-dimensional (3D) model based on threading and ab initio techniques. The top-ranked model was selected based on the C-score (confidence score) and subjected to further refinement to improve its stereo chemical quality and resolve structural inaccuracies, particularly in flexible loop regions. The initial model generated by I-TASSER had unresolved loop regions, which were refined using ModLoop and Galaxy Refine web server tools [32]. The ModLoop tool was employed to adjust the flexible regions of the structure, particularly those between the NS2B cofactor and the NS3 catalytic domain. Subsequently, the Galaxy Refine web server optimized the overall geometry by minimizing steric clashes and improving the side-chain conformations [33]. This step enhanced the model’s reliability, with improved Ramachandran plot statistics and reduced energy scores [34]. The final refined structure was validated using PROCHECK and Verify3D to ensure its quality. A structural comparison between the refined model and the template structure (PDB ID: 2FOM) indicated that the overall structure of the Dengue NS2B– NS3 protease remained conserved. The calculated backbone RMSD value of 2.757 Å suggests that moderate conformational differences was there during the refinement process. Most of these deviations were observed in flexible loop, surface-exposed regions whereas the core secondary structural elements and catalytic residues retained stable conformations. The final structural model of Dengue NS2B-NS3 protease (***Fig.5***) revealed a compact architecture, with the NS3 domain forming a classic protease fold consisting of β-strands and α-helices. The NS2B cofactor was positioned adjacent to the catalytic triad to stabilizing. The structural features observed are consistent with previous studies on homologous *flavivirus* proteases, reinforcing the biological relevance of the model.

**Fig. 5.**
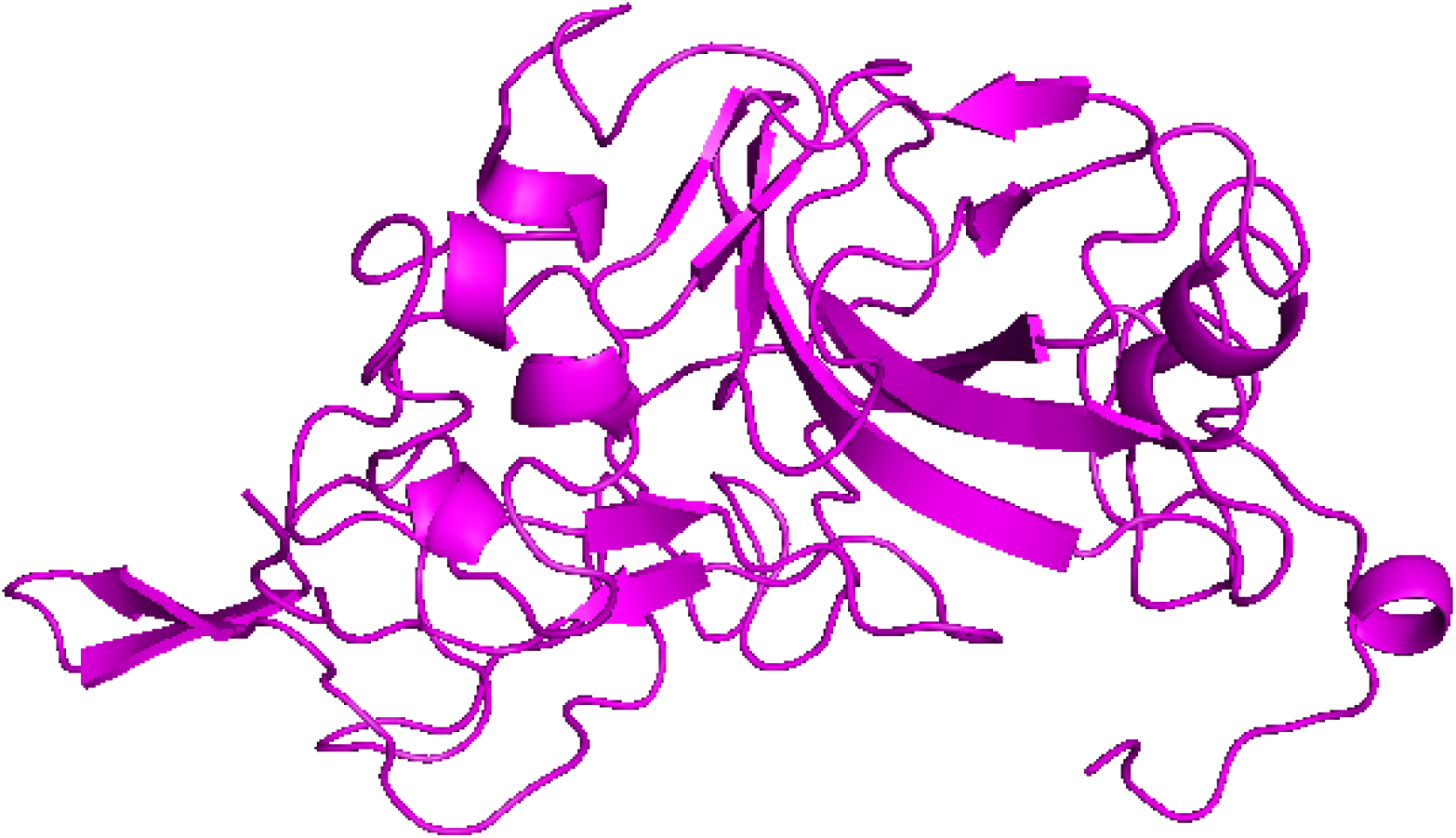
Modelled structure of Dengue NS2B-NS3 protease. The 3D structure was predicted using I-TASSER and refined with ModLoop and GalaxyRefine.

### MD Simulations

The molecular dynamics simulations conducted on the Dengue NS2B-NS3 protein under different pH conditions reveal critical insights into its structural stability and dynamic behavior. The analysis includes Root Mean Square Fluctuation (RMSF), Root Mean Square Deviation (RMSD), and hydrogen bond distribution. These parameters provide a comprehensive understanding of how pH influences the conformational stability of the protein, which is crucial for its enzymatic activity and potential therapeutic targeting.

### RMSD Analysis

The RMSD which measures global structural deviations, further corroborates the pH-dependent stability of Dengue NS2B-NS3pro. At pH 4, the RMSD remained relatively stable around 0.5 – 0.6 nm throughout the simulation, indicating minimal structural deviation and a highly stable conformation under acidic conditions. At pH 5 and 6, similar trends were observed, with RMSD values fluctuating slightly but remaining below 0.8 nm, suggesting good stability in mildly acidic environments. At pH 7.5, the RMSD stabilizes around 0.5 nm throughout the simulation, indicating the maintenance of the protein’s overall structure **(*Fig.6*).** This stability reflects the protein’s natural conformational preference under physiological conditions. In, under acidic (pH 4 and 5) and mildly acidic (pH 6) conditions, the RMSD values are slightly elevated and suggesting minor structural perturbations. In contrast, at basic condition, the RMSD values rise significantly, exceeding 1.5 nm in the case of pH 8. This suggests that the protein undergoes substantial conformational changes under alkaline conditions, potentially impacting its enzymatic activity. The higher RMSD at pH 8 aligns with the increased flexibility observed in the RMSF analysis. The increased flexibility observed at pH 8 and 9 may reflect conformational adjustments needed for activity under mildly alkaline conditions. However, the significant instability at pH 10 highlights the protein’s intolerance to highly alkaline environments, likely due to disruption of critical intramolecular interactions. Similar findings have been reported in studies examining the pH-dependent dynamics of other viral proteases, where extreme pH conditions disrupt the structural stability required for enzymatic function [35].

**Fig. 6.**
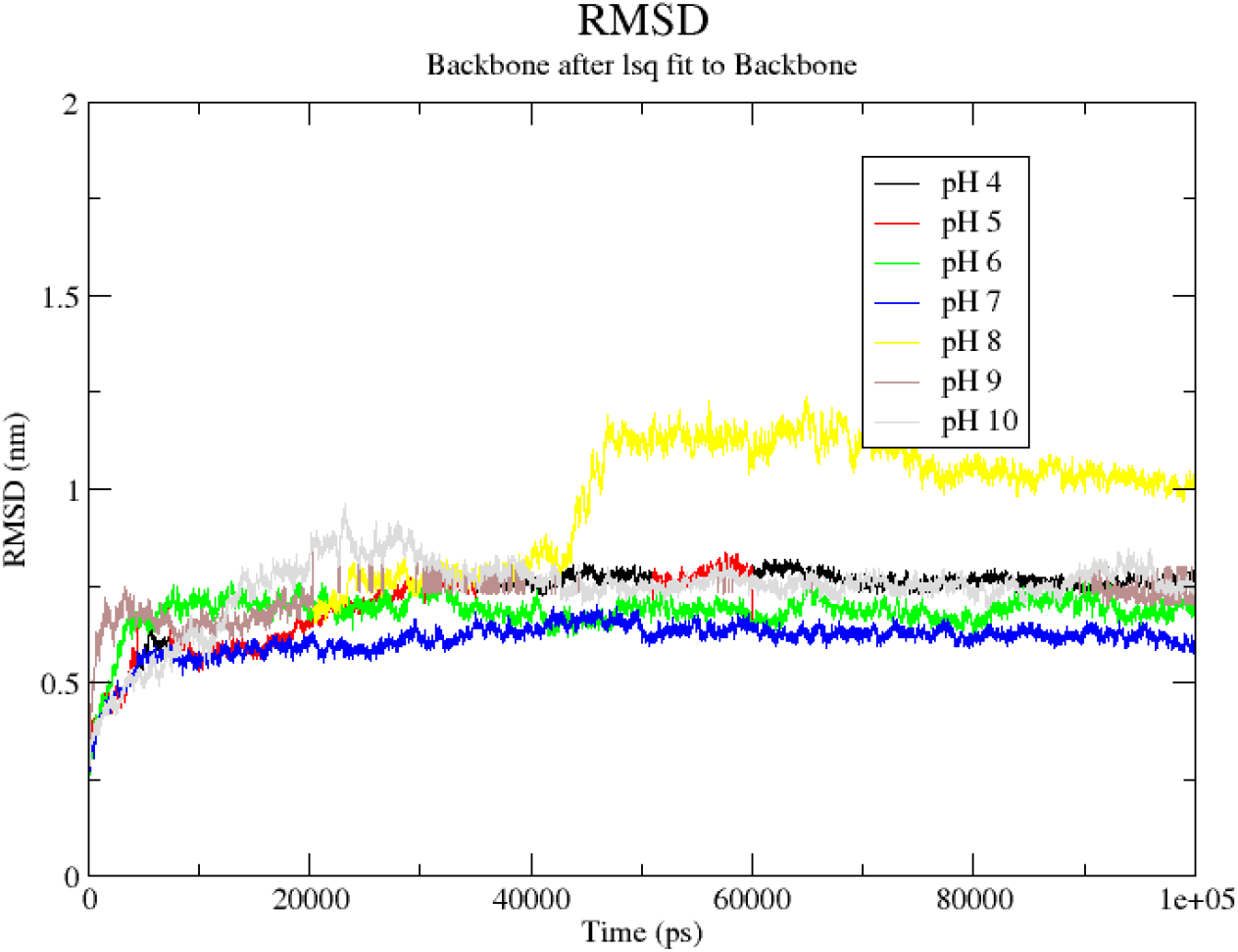
The RMSD figure highlights the protein’s dynamic behavior and structural stability across acidic and neutral environments during the 100 ns simulation.

### RMSF Analysis

The RMSF (Root Mean Square Fluctuation) analysis of Dengue NS2b-NS3 protein across various pH conditions (4, 5, 6, 7, 8, 9, and 10) during 100 ns molecular dynamics (MD) simulations is shown in the graph. The fluctuations provide residue-level insights into the protein’s flexibility and dynamic stability under different pH conditions. At acidic pH value 4 and 5, the RMSF values are relatively low for most residues, averaging between 0.2 – 0.6 nm. This indicates that the protein backbone remains stable with limited fluctuations in these conditions. At neutral pH 7.5, the RMSF values remain consistently low, further confirming the structural stability of the protein under physiological conditions. At pH 8 and 9, certain residues show increased flexibility, with RMSF values rising to approximately 1.0 nm. This suggests moderate destabilization in these mildly alkaline environments. At pH 10, significant peaks are observed, particularly for residues around positions 90 – 120, where RMSF values exceed 1.5 nm. This indicates localized destabilization or structural rearrangements, likely due to the disruption of hydrogen bonding or electrostatic interactions at high pH levels. Residues 50–90 exhibit minimal fluctuations across all pH conditions, indicating their involvement in a stable core structure. Residues around positions 100 – 130 exhibit the highest flexibility at pH 10, likely corresponding to loop or surface-exposed regions more prone to pH-dependent conformational changes. Terminal regions also show slightly higher fluctuations, consistent with their inherent flexibility in protein structures. The RMSF analysis demonstrates that Dengue NS2b-NS3 maintains structural stability under acidic (pH 4 – 6) and neutral (pH 7.5) conditions, as reflected by the low RMSF values across most residues. This stability is critical for the protease’s functionality under physiological and intracellular conditions, where slight acidic to neutral pH predominates. However, at higher pH values (pH 8 – 10), the protein displays increased flexibility, with notable fluctuations in specific regions such as residues 90–120 (***Fig.7*.**). This region may correspond to loop or active site residues undergoing conformational changes due to altered protonation states at higher pH, affecting intramolecular interactions. The significant fluctuations at pH 10 suggest partial destabilization or unfolding, which could impair enzymatic activity and structural integrity.

**Fig. 7.**
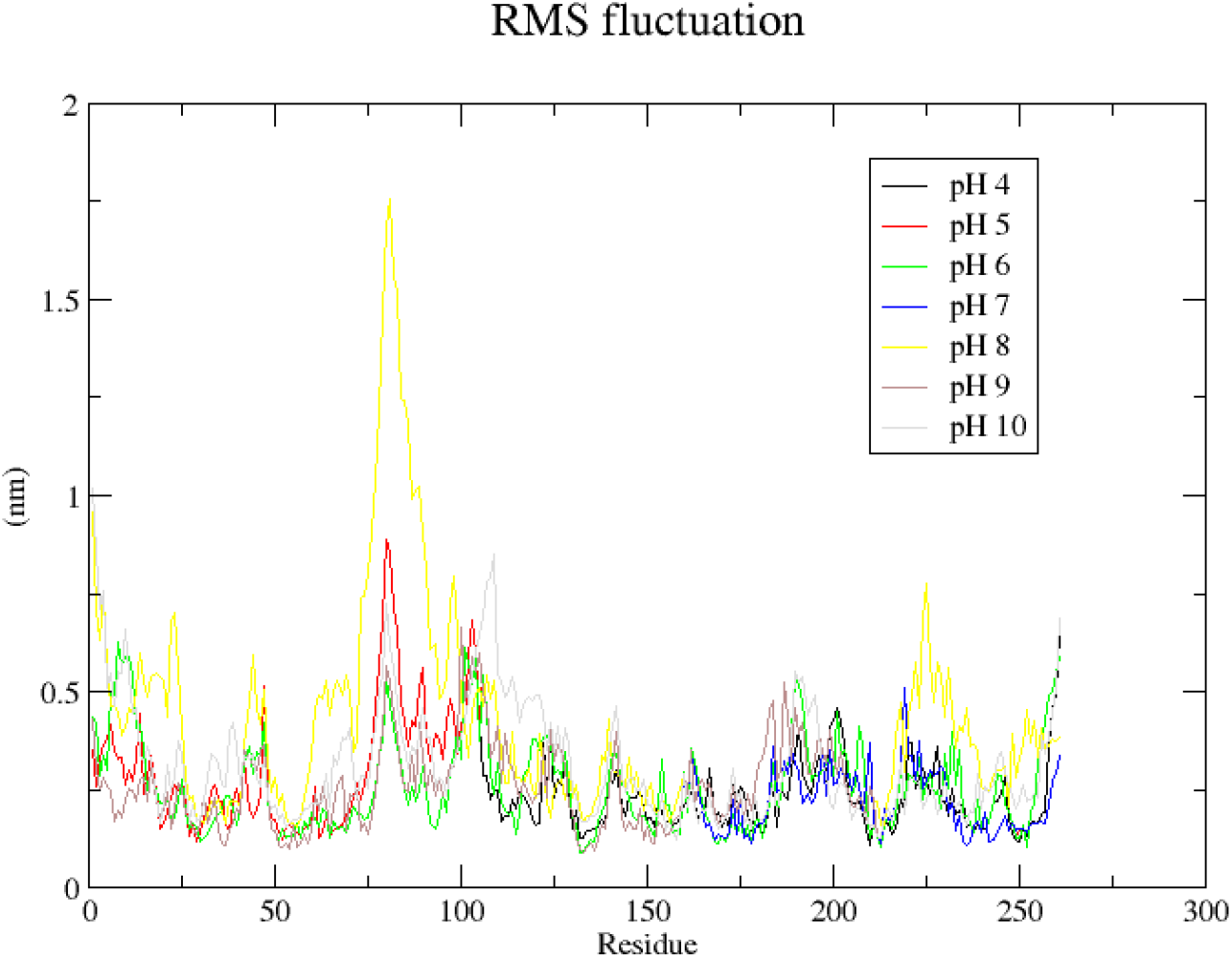
Depicts the RMSF (Root Mean Square Fluctuation) graph from MD simulations of the Dengue NS2b-NS3 protein under various pH conditions across acidic, neutral, and alkaline environments.

The peaks in the RMSF graph likely correspond to flexible loop regions, which are inherently more dynamic and responsive to environmental conditions [36,37]. These findings suggest that the protein retains its functional conformation at neutral pH while becoming more flexible and potentially less efficient at pH extremes.

### Hydrogen Bond Analysis

Hydrogen bonds play a critical role in maintaining the structural stability of proteins, and their distribution under different pH conditions provides valuable insights. At pH 4, the protein maintained a consistent number of hydrogen bonds (120–140), indicating stable intramolecular interactions. A similar trend was observed at pH 5, with a slight increase in hydrogen bond count, reaching up to 150. At pH 6, the number of hydrogen bonds remained stable, fluctuating between 130 and 140, suggesting good structural integrity under mildly acidic conditions. At pH 7.5, the hydrogen bond count remained steady between 120 and 140, highlighting the protein’s optimal stability at physiological conditions. The data corroborate the expected structural stability of the Dengue NS2b-NS3 protein at neutral pH. At pH 8, the hydrogen bond count exhibited larger fluctuations, ranging from 140 to 160, reflecting moderate changes in structural stability. At pH 9, the number of hydrogen bonds increased significantly, peaking at approximately 180. This suggests enhanced hydrogen bonding, potentially due to altered intramolecular interactions caused by the alkaline environment. At pH 10, the hydrogen bond count displayed considerable fluctuations (160 – 200), indicating structural destabilization and possible unfolding under highly alkaline conditions (***Fig.8***). These results highlight the importance of maintaining a balanced pH for the protein’s stability, as extreme pH conditions lead to structural disruptions

**Fig. 8.**
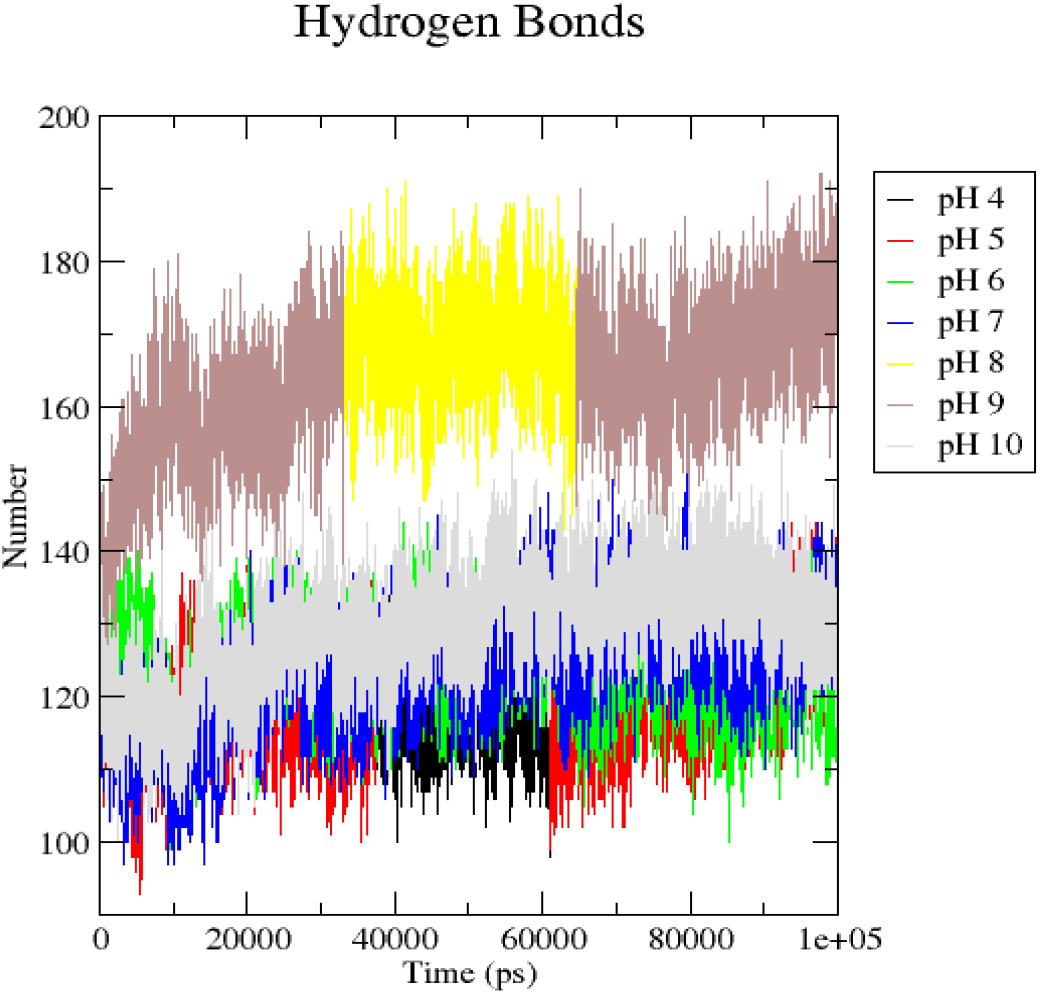
Illustrates the hydrogen bond graph for the Dengue NS2b-NS3 protein at various pH conditions.

The Dengue NS2b-NS3 protein shows consistent hydrogen bond counts under acidic to neutral conditions (pH 4 – 7), indicating stable intramolecular interactions and maintenance of its native conformation essential for enzymatic activity [22]. At alkaline (pH 8 – 10), increased and fluctuating hydrogen bonds suggest a dynamic structural response. While pH 9 exhibits transient stabilization. And pH 10 shows significant fluctuations, indicating partial unfolding and compromised stability. These results align with reports that extreme pH disrupts protein structure [41]. . The MD simulation results showed trends similar to those observed in the CD spectroscopy experiments. Under neutral to mildly alkaline conditions (pH 7.5–8), the protein displayed lower RMSD and RMSF values along with stable hydrogen bonding patterns, indicating better structural stability. These findings were consistent with the CD spectra, which showed preservation of secondary structural elements under the same conditions. In contrast, acidic and highly alkaline environments resulted in increased structural fluctuations and altered hydrogen bond networks during the simulations, correlating with the partial unfolding and destabilization observed experimentally in the CD analysis.

### DSSP

The DSSP analysis of the Dengue NS2B-NS3 protease across a range of pH conditions (pH 4, 5, 6, 7, 8, 9 and 10) reveals distinct secondary structure transitions, emphasizing the protease’s pH-dependent stability and its correlation with enzymatic activity. The DSSP plot at pH 4 (highly acidic condition) indicates substantial secondary structure loss due to extensive denaturation (***Fig.9***). Within the first 30 ns, partial α-helices and β-sheets remain stable, but residues 150–170 undergo significant transitions to random coils (green regions), indicating early destabilization [10]. By 30 – 40 ns, the α-helices in residues 120 – 140 and β-sheets in residues 180 – 200 are disrupted, with the structure becoming predominantly unstructured. From 50 – 70 ns, residues 50 – 100 transitions to random coils, suggesting complete unfolding of the loop regions (***Fig.10*.**).

**Fig. 9.**
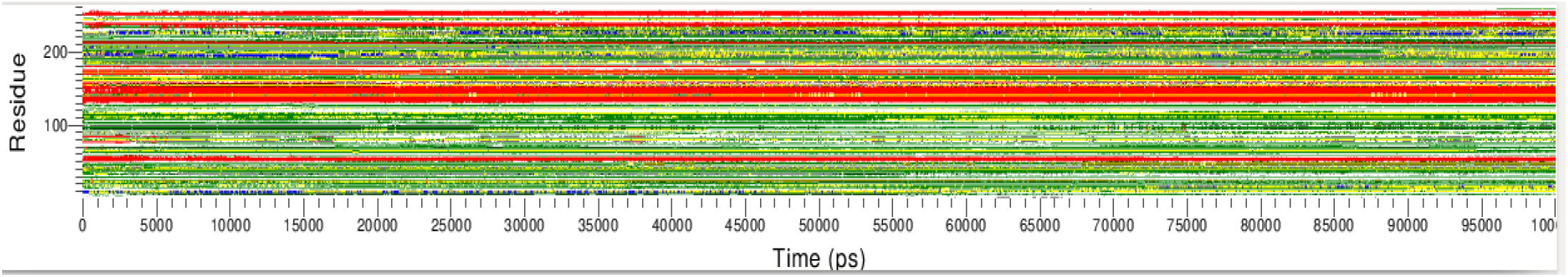
DSSP analysis of Dengue NS2B-NS3 protease at pH 4. The plot indicates minimal secondary structure transitions, with evident denaturation of α-helices and β-sheets over 100 ns.

**Fig. 10.**
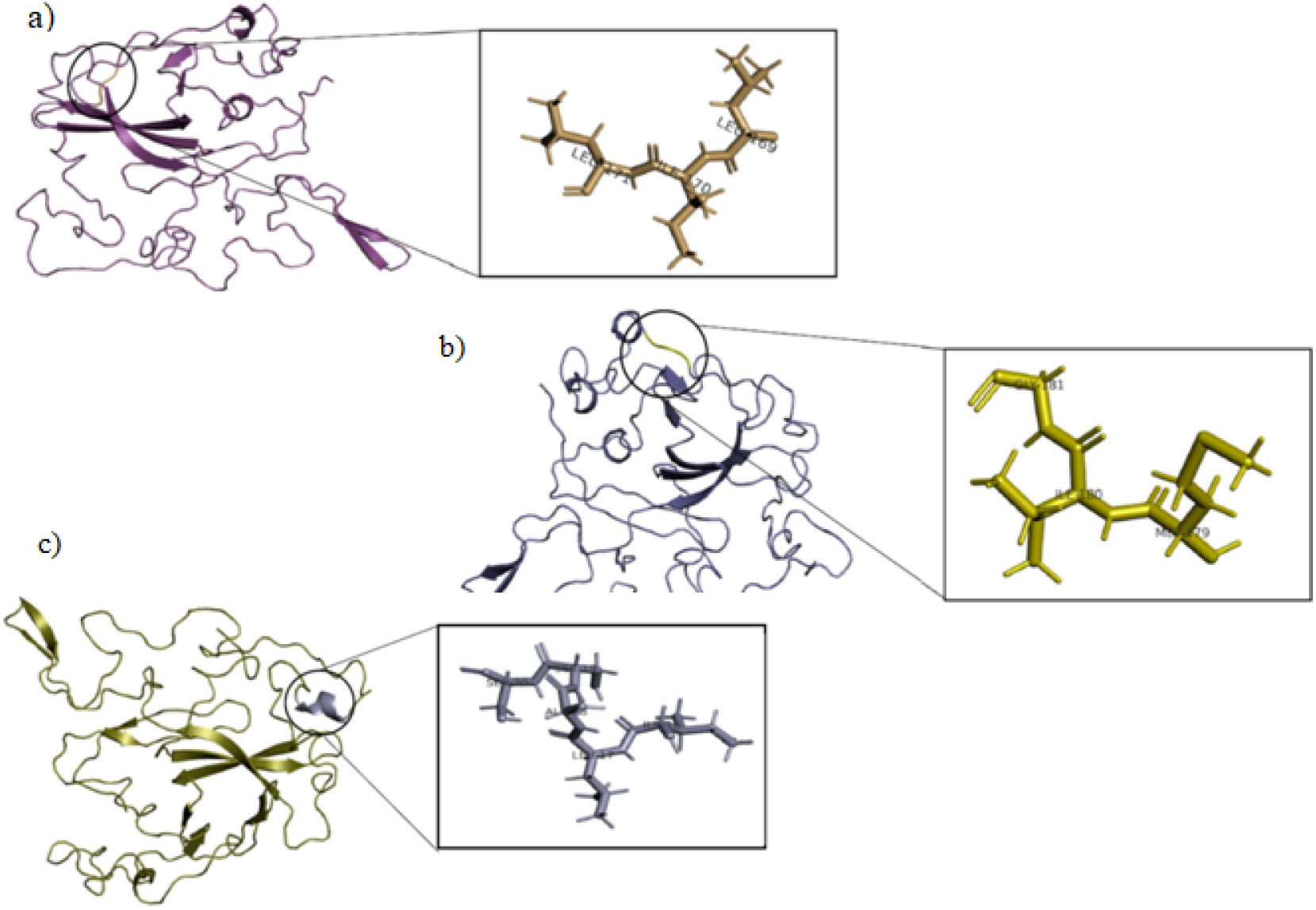
The plot highlights significant secondary structure transitions, with progressive denaturation of α-helices and β-sheets, resulting in a predominantly unstructured conformation over time. a) 169 - 171 amino acid changes from beta sheet to loop at 35ns. b) 179 - 181 amino acid residue changes from beta sheet to loop at 60 ns. c) 16 - 20 amino acid residues change from loop to alpha helix at 75ns.

By the end of the simulation (70 – 100 ns), the protein is largely denatured, with minimal β-sheet content observed in residues 10 – 30. The secondary structure at pH 5 undergoes noticeable but less severe changes rather at pH 4, highlighting partial preservation of the native conformation. During the initial 20 ns, α-helices and β-sheets, particularly in residues 50 – 200, remain mostly intact, similar to pH 6. However, slight destabilization is observed in loop regions (residues 80 – 100), which transition transiently into random coils. From 50-70 ns, residues 50-100 exhibit moderate transitions to random coil conformations, indicating progressive destabilization of loop regions .

Compared to pH 4, where the structure is largely unstructured by this point, pH 5 retains significant β-sheet and α-helical content (***Fig.11***). Between 70 – 100 ns, α-helices in residues 120 – 140 and β-sheets in residues 180 – 200 shows increased disruptions, yet the overall structure at pH 5 remains more ordered than at pH 4, with noticeable retention of secondary structure in residues 10 – 30 and 150–170 (***Fig.12***).

**Fig. 11.**
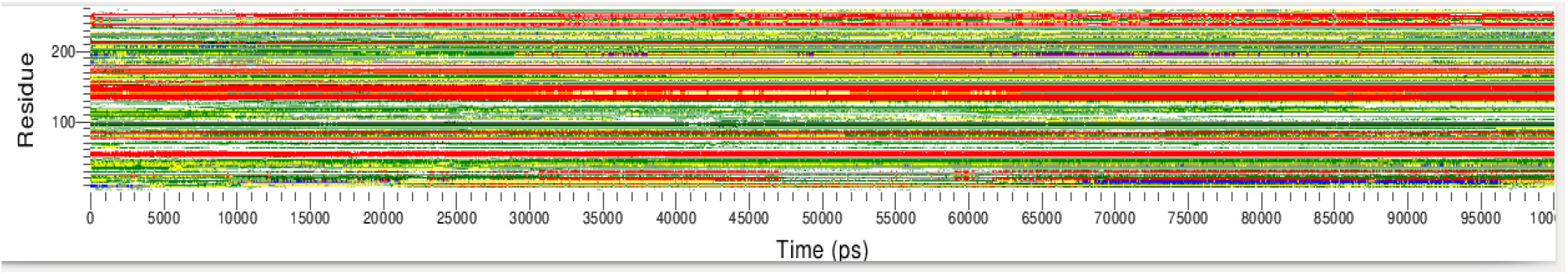
DSSP analysis of Dengue NS2B-NS3 protease at pH 5. The plot indicates minimal secondary structure transitions, with evident denaturation of α-helices and β-sheets over

**Fig. 12.**
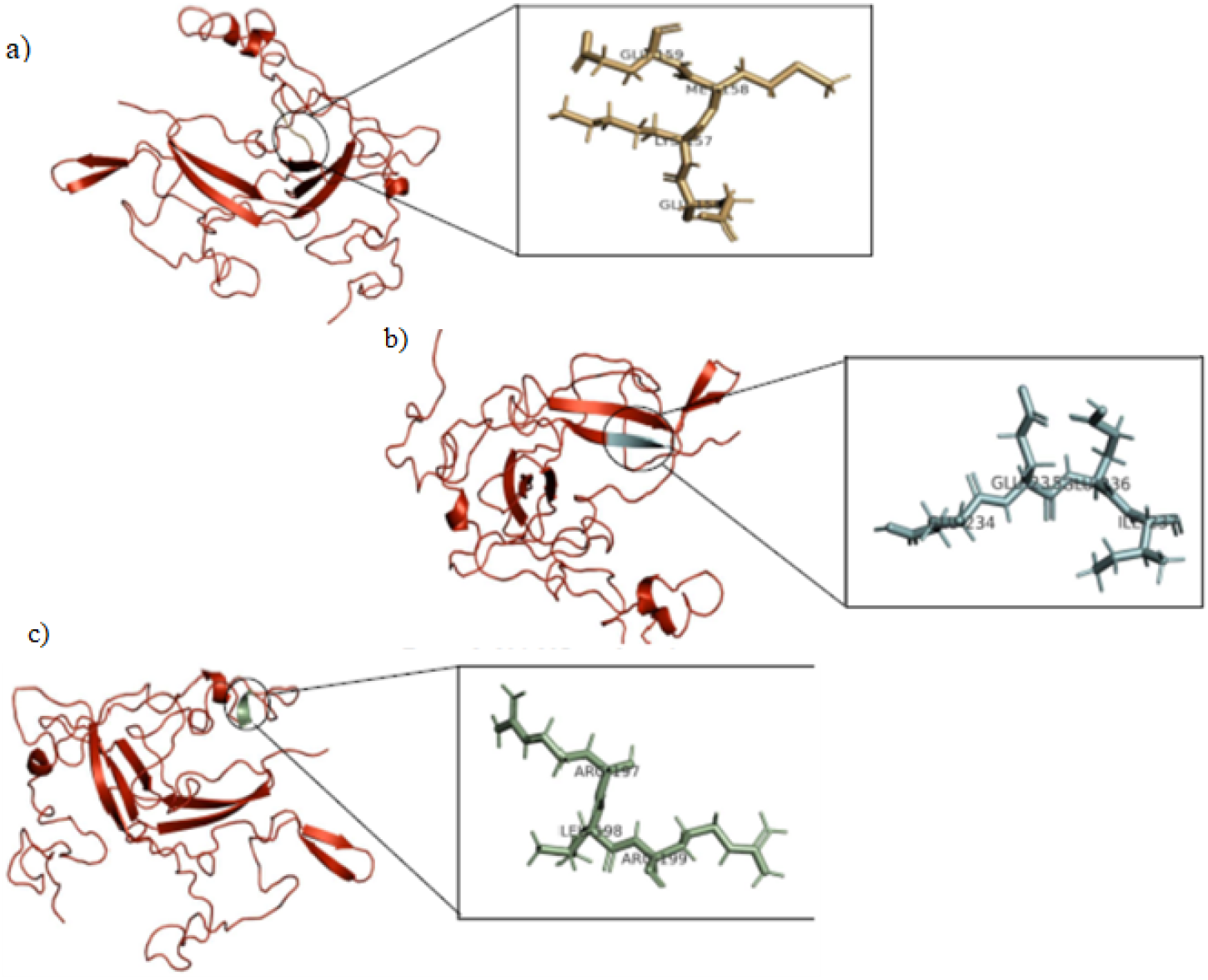
The plot highlights significant secondary structure transitions, with progressive denaturation of α-helices and β-sheets, resulting in a predominantly unstructured conformation over time. a) 68 - 70 amino acid changes from beta sheet to loop at 55ns. b) 78 - 80 amino acid residue changes from loop to sheet at 65ns. c) 127 - 131 amino acid residues change from loop to alpha helix at 73ns.

The DSSP plot at pH 6 (mildly acidic condition) shows moderate structural perturbations compared to the more acidic pH 4. During the initial 20 ns, α-helices and β-sheets dominate, particularly in residues 50 – 200 [38]. Residues 80 – 100 shows transient shifts to random coils. At 40 – 50 ns, α-helices in residues 120 – 140 partially unwind into random coils, and residues 180 – 200 shows disruptions in β-sheets (***Fig.13***).

**Fig. 13.**
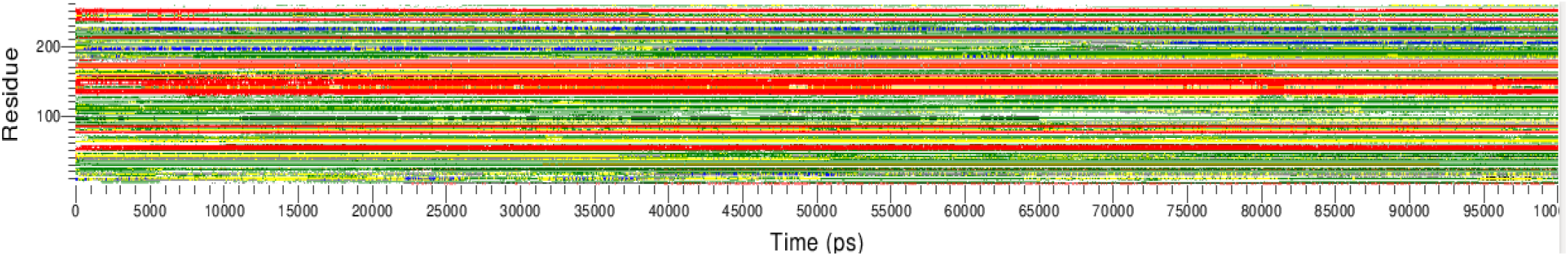
DSSP analysis of Dengue NS2B-NS3 protease at pH 6. The plot indicates minimal secondary structure transitions, protease shows partial destabilization over 100 ns

Between 60 – 80 ns, there are localized β-sheet disruptions due to protonation effects. By the end of the simulation, residues 50 – 70 transitions to unstructured conformations, indicating mild but progressive destabilization (***Fig.14***).

**Fig. 14.**
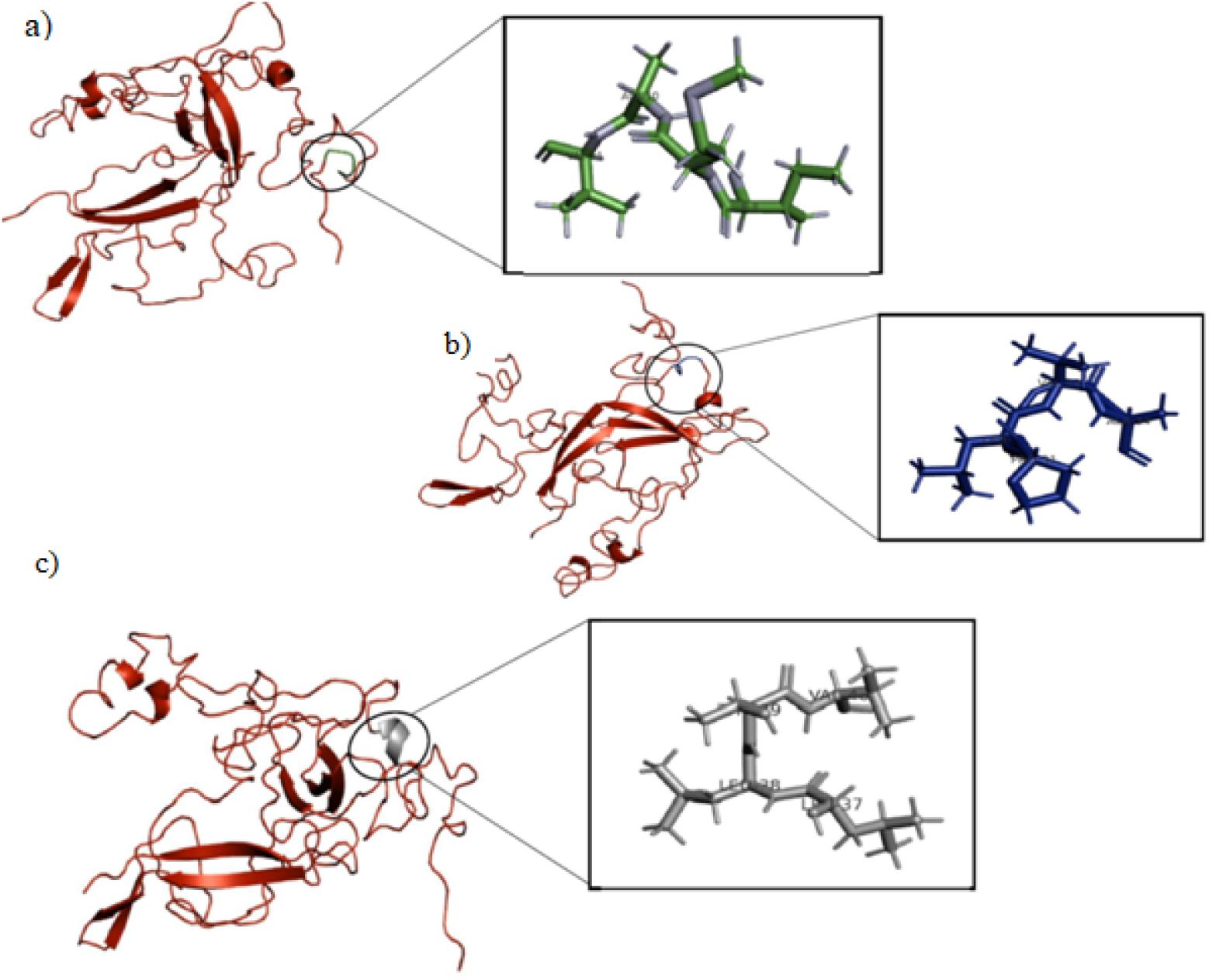
DSSP analysis of Dengue NS2B-NS3 protease at pH 6. a) 68 - 70 amino acid changes from beta sheet to loop at 45ns. b) 78 - 80 amino acid residue changes from sheet to loop at 55ns. c) 127 - 131 amino acid residues change from loop to alpha helix at 80ns.

The protease structure is highly stable at pH 7.5 (Neutral condition), as shown in the DSSP plot [39]. From 0 – 20 ns, α-helices (red) and β-sheets (yellow) are well-maintained. Only minor random coil formation is observed in flexible loop regions (residues 50 – 70). Throughout the simulation, the catalytic core and structural domains (residues 120 – 200) exhibit minimal transitions, indicating robust stability (***Fig.15***). Between 50 – 100 ns, loop flexibility increases, but the overall secondary structure remains intact (***Fig.16***).

**Fig. 15.**
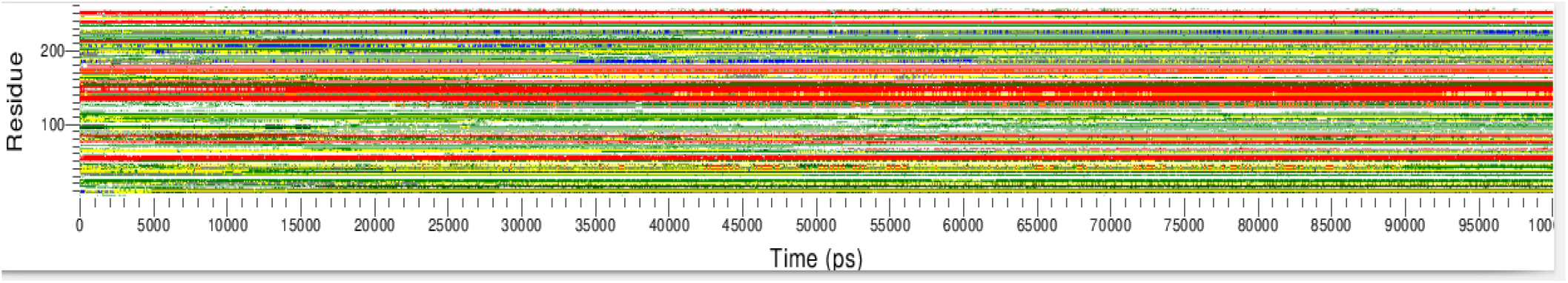
DSSP analysis of Dengue NS2B-NS3 protease at pH 7.5. The plot indicates maximal stability over 100 ns.

**Fig. 16.**
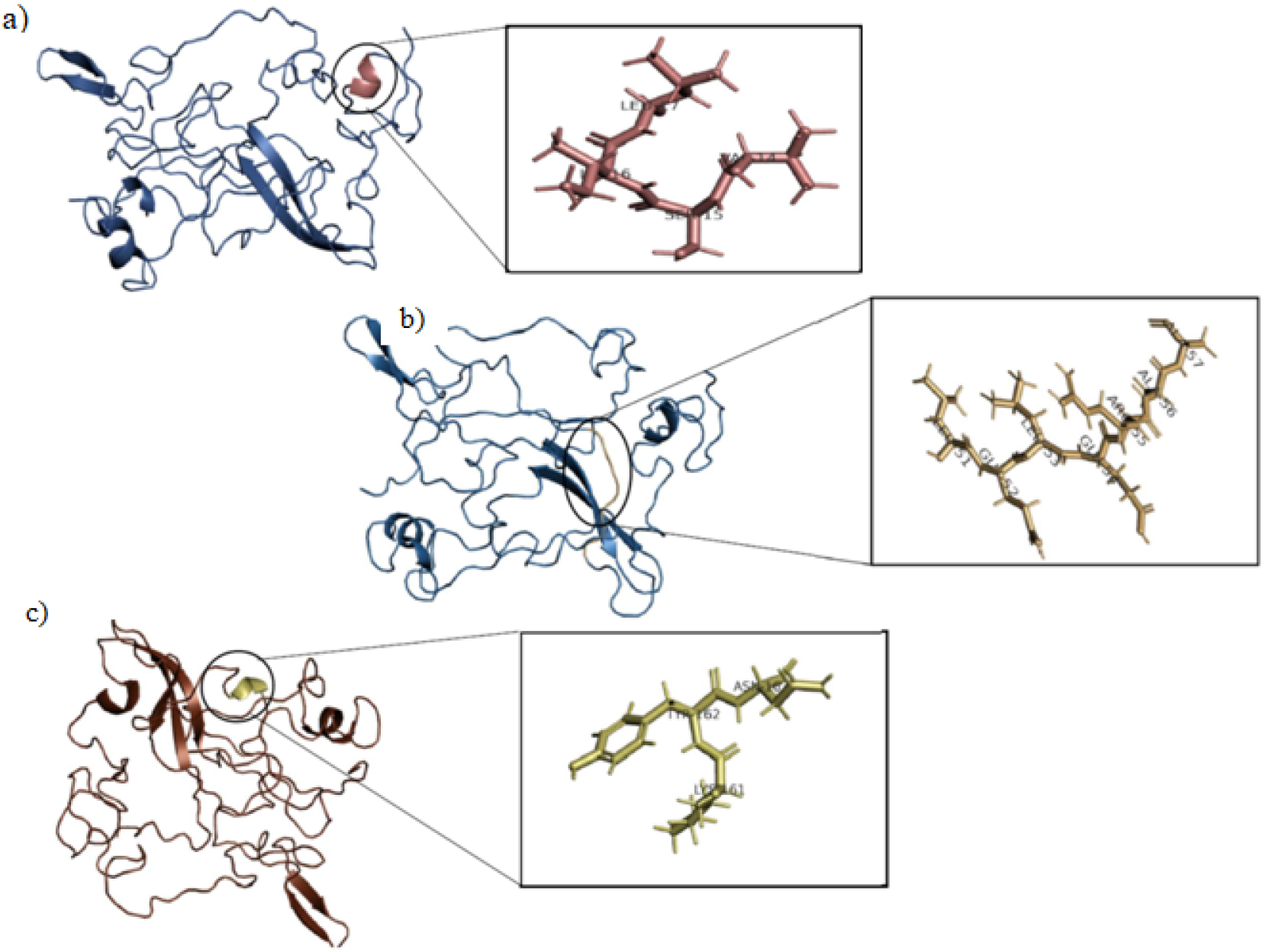
DSSP analysis of Dengue NS2B-NS3 protease at pH 7.5. a) 64 - 69 amino acid changes from loop to helix at 53ns. b) 58-62 amino acid residue changes from sheet to loop at 55ns. c) 121 - 125 amino acid residues change from loop to alpha helix at 75ns.

At pH 8 (mildly alkaline condition), the protease maintains significant structural integrity with minor transitions observed in flexible regions (***Fig.17***). In the initial 20 ns, stable α-helices and β-sheets are observed, particularly in the catalytic triad and the NS2B cofactor-binding domain. Between 20 – 60 ns, the central β-sheet (residues 120 – 160) remains stable, with transient transitions of α-helices (residues 50 – 60) to 3 - 10 helices at around 35 ns. By 60 - 100 ns, flexibility in the NS2B domain (residues 80 – 90) leads to brief coil transitions (***Fig.18***), but the structure largely retains its integrity.

**Fig. 17.**
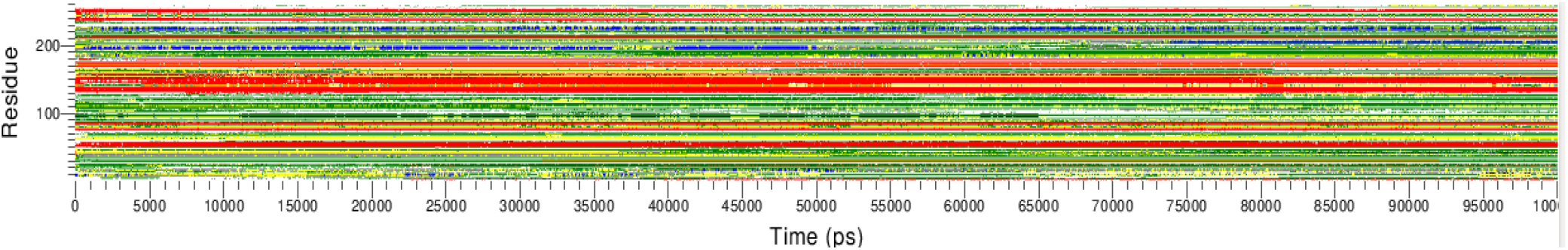
DSSP analysis of Dengue NS2B-NS3 protease at pH 8. The plot indicates minimal secondary structure transitions with stable stable α-helices and β-sheets over 100 ns.

**Fig. 18.**
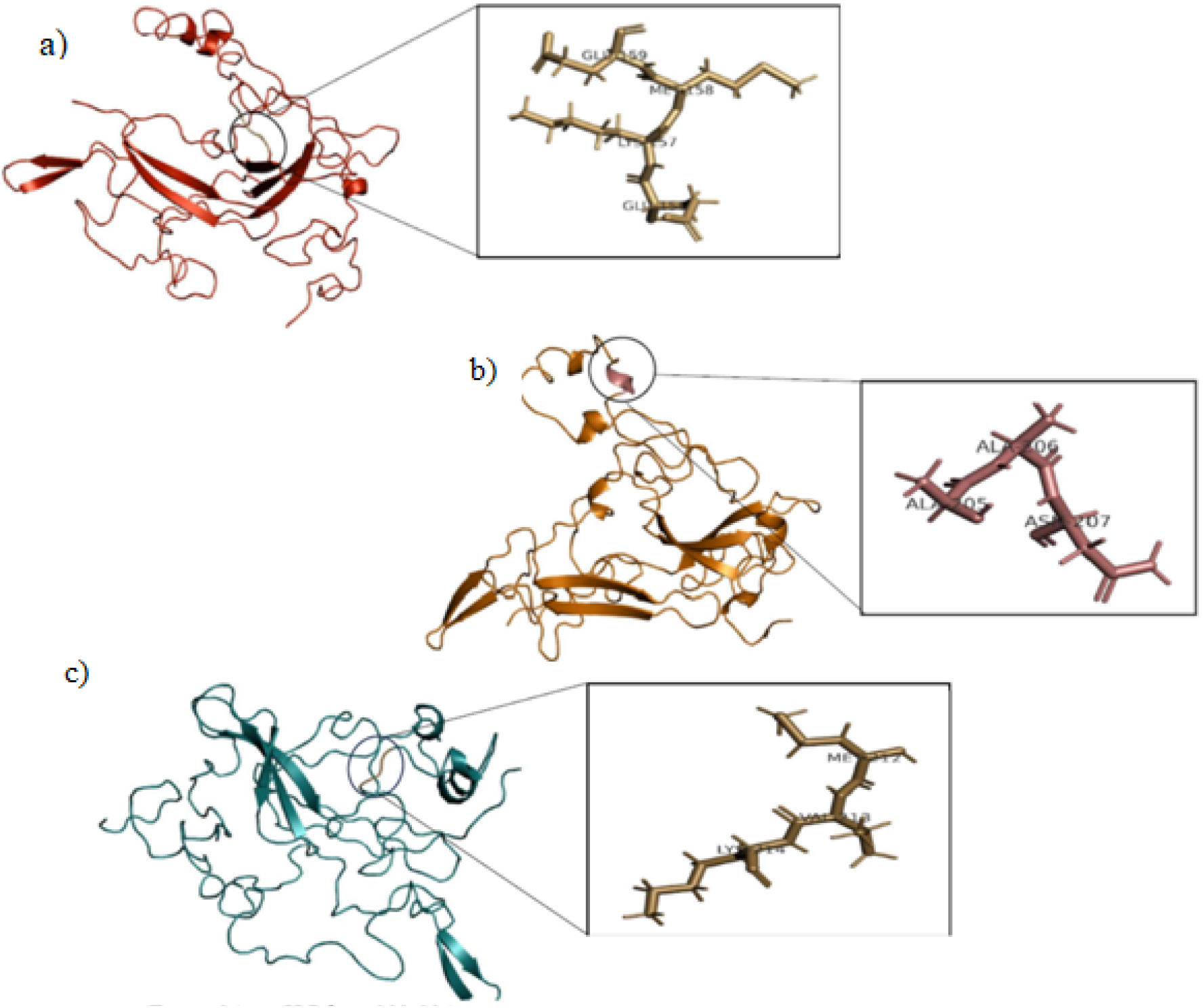
DSSP analysis of Dengue NS2B-NS3 protease at pH 8. a) 84 - 90 amino acid changes from sheet to loop at 60ns. b) 58 - 62 amino acid residue changes from loop to helix at 35ns. c) 78 - 82 amino acid residues change from loop to alpha helix at 63ns

At pH 9 (alkaline solution) it is likely that the first 20 ns, the protease retains most of its secondary structure (***Fig.19***), However, between 20 – 60 ns, the central β-sheet (residues 130 - 50) might experience partial hydrogen bond loss, and loop regions (residues 60–80) could transition into random coils. By 60 – 100 ns, the NS2B domain (residues 80 – 100) might shows moderate disorder, with some β-sheet disruptions in the core region. Overall, pH 9 is expected to result in a gradual destabilization (***Fig.20***).

**Fig. 19.**
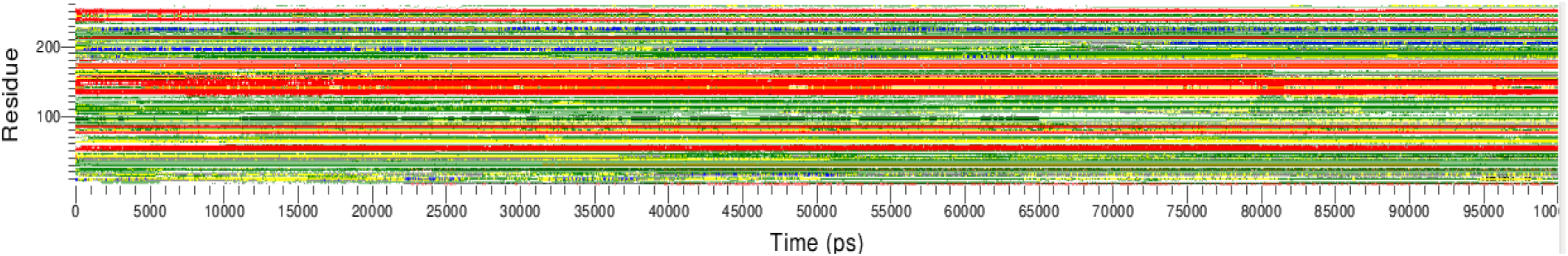
DSSP analysis of Dengue NS2B-NS3 protease at pH 9. The plot indicates minimal secondary structure transitions, with stable α-helices and β-sheets over 100 ns

**Fig. 20.**
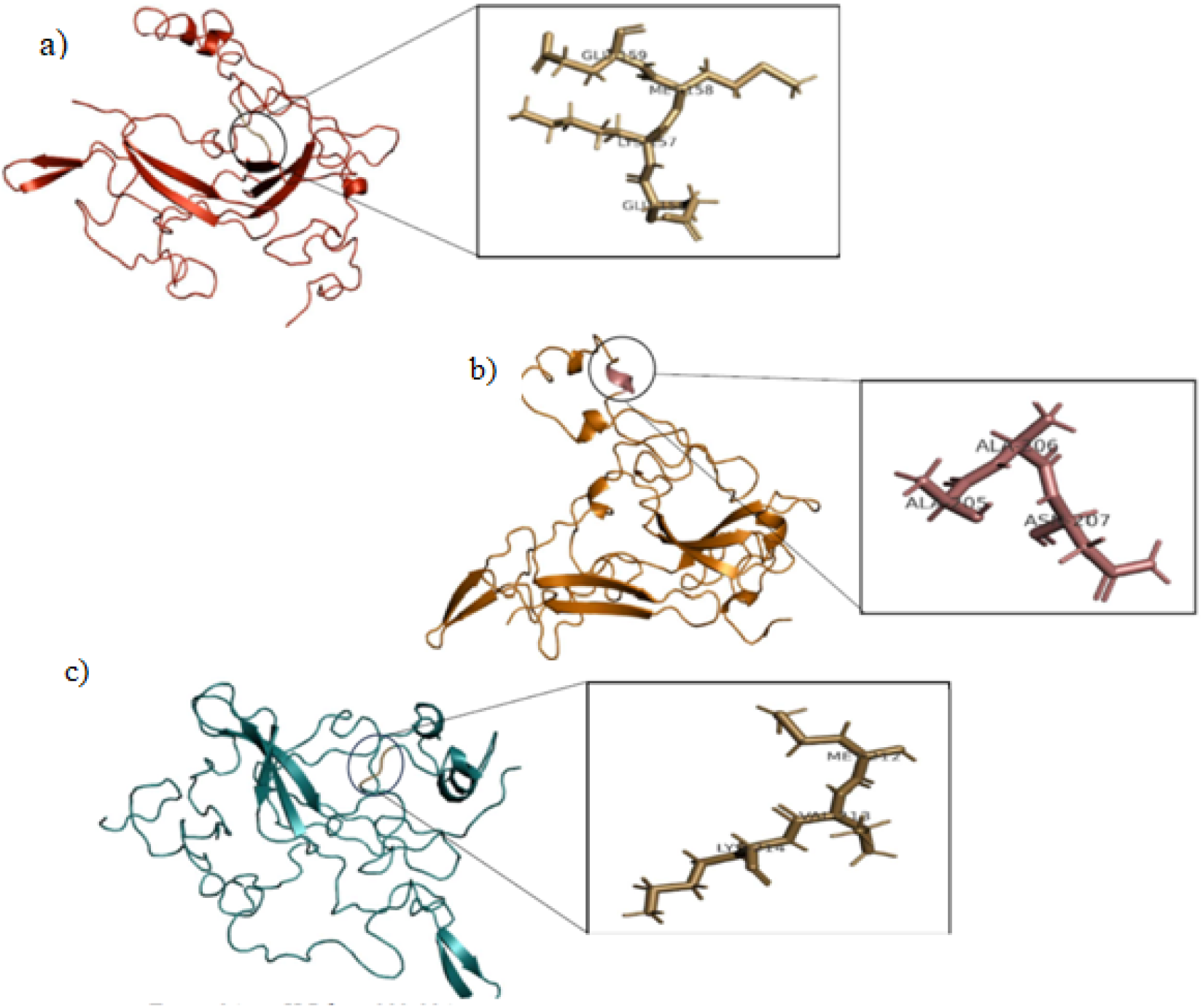
DSSP analysis of Dengue NS2B-NS3 protease at pH 9. a) 135 - 139 amino acid changes from sheet to loop at 60ns. b) 89 - 94 amino acid residue changes from loop to helix at 85ns. c) 68 - 73 amino acid residue changes from helix to loop at 50ns

Elevated pH 10 (highly alkaline condition) induces substantial structural alterations, resulting in partial destabilization of the protease. By 20 ns, helices near the catalytic triad (residues 45 – 65) shows instability (***Fig.21***), with transitions into coils. Between 20 – 60 ns, β-sheets in the core region (residues 120 – 160) partially lose hydrogen bonding, while loop regions (residues 60 – 80) transition to coils. From 60 – 100 ns, residues in the NS2B domain (80 – 100) are fully disordered, and the core β-sheets exhibit significant destabilization (***Fig.22***). The overall structure retains less than 70% of its original secondary structure.

**Fig. 21.**
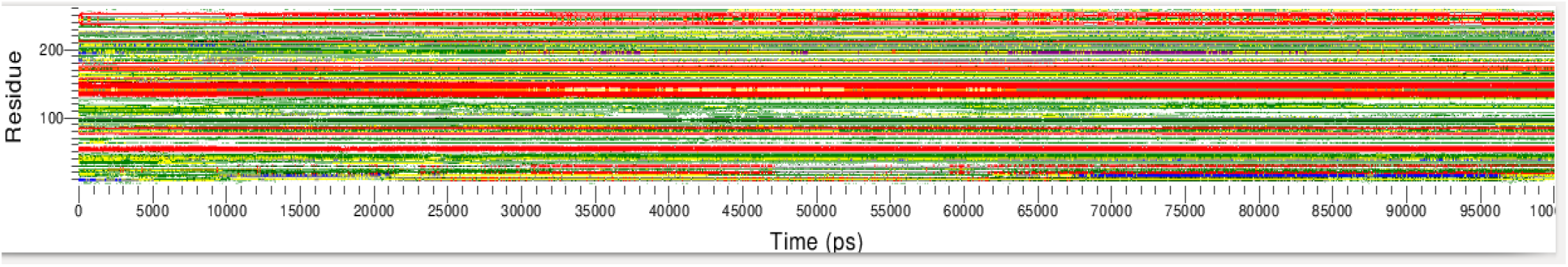
DSSP analysis of Dengue NS2B-NS3 protease at pH 10. The plot shows significant secondary structure loss, including unwinding of helices and destabilization of β-sheets, particularly after 60 ns.

**Fig. 22.**
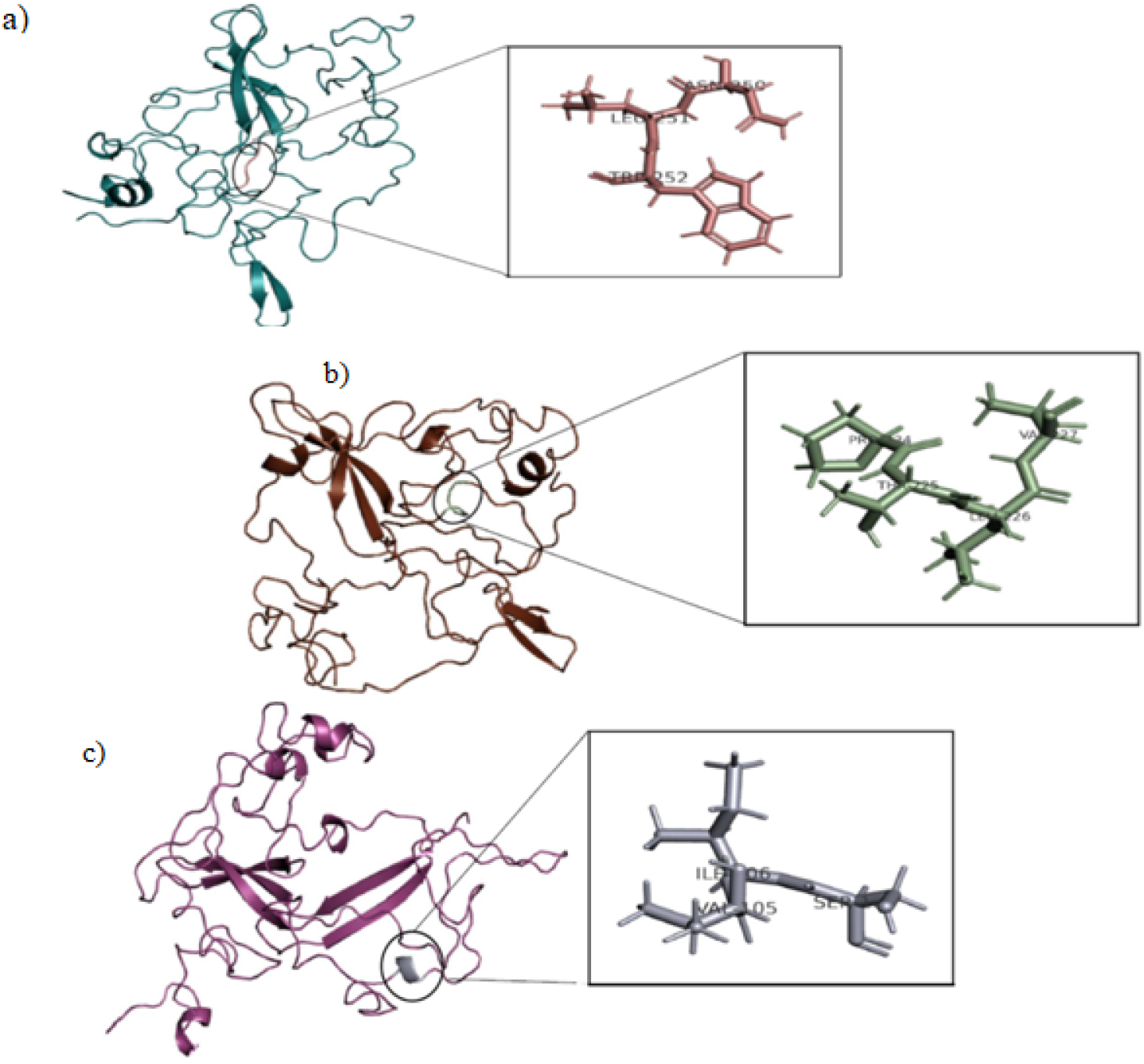
DSSP analysis of Dengue NS2B-NS3 protease at pH 10. a) 58 - 65 amino acid changes from sheet to loop a 20ns. b) 60 - 65 amino acid residues change from sheet to loop at 70ns. c) 89 - 94 amino acid residue changes from loop to helix at 90ns.

The secondary structural dynamics of the Dengue NS2B-NS3 protease across pH conditions highlight its sensitivity to pH changes, which directly affect its structural integrity and enzymatic function. At pH 4, extensive denaturation is evident, with loss of α-helices and β-sheets. This is likely due to protonation of acidic residues (e.g., Glu, Asp) and disruption of critical salt bridges, which destabilize the hydrogen bond network. At pH 5 demonstrate that the protein exhibits a mixture of monomeric and aggregated states with a significant degree of polydispersity, likely due to the influence of acidic conditions on structural stability. This analysis underscores the effect of low pH on protein aggregation behavior, with the acidic environment potentially leading to destabilization and aggregation of the protein structure. At pH 6, the protease shows partial destabilization, particularly in loop and helix regions. While the structure is less stable compared to neutral pH, it retains substantial secondary structure, suggesting that mild acidity facilitates flexibility necessary for conformational changes during viral processes. At pH 7.5, the protease displays maximal structural stability, with α-helices and β-sheets well-preserved throughout the simulation. The intrinsic flexibility of loop regions correlates with substrate binding and catalytic activity, consistent with physiological conditions. At pH 8, the protease retains over 90% of its secondary structure, with minor coil transitions in flexible regions, indicating adaptability and robustness. The aggregation behavior observed at pH 9 may have implications for the protein’s structural and functional studies. At pH 10, significant disruptions are observed, including unwinding of helices near the active site and disordered loops. The destabilization of β-sheets compromises the catalytic core, suggesting reduced enzymatic activity under alkaline conditions.

The comparison of CD spectroscopy and DSSP analysis reveals consistent structural changes in the Dengue NS2B-NS3 protease across different pH conditions, though with varying levels of detail. At pH 4, both method shows significant destabilization, with CD indicating high random coil content (37.9%) and decreased helical content (27.6%), suggesting partial unfolding. DSSP corroborates this with transitions to random coils and loss of secondary structure. At pH 5, the CD results indicate the secondary structure composition as 25.1% α-helix, 5.3% β-sheet, 34.5% turns, and 37.9% random coils, reflecting a partial preservation of the protein’s native conformation. DSSP analysis supports this by showing that α-helices and β-sheets, particularly in residues 50–200, remain mostly intact during the initial 20 ns, similar to the behavior at pH 6. At pH 6, the protease exhibits partial destabilization in both analyses, with CD showing 28.4% helical content and 37.9% random coils, while DSSP reveals transient coil transitions and some unwinding of helices, though more structure is retained compared to pH 4. At pH 7.5, both CD and DSSP indicate maximal stability, with CD showing a well-balanced secondary structure and DSSP confirming stable α-helices and β-sheets. At pH 8, CD shows a slight shift towards random coils (47.3%), indicating some structural reorganization, which is also reflected in DSSP with increased flexibility in the NS2B domain. At pH 9, the CD analysis shows a secondary structure composition of 31% α-helix, 0% β-sheet, 30.6% turns and 38.4% random coils, indicating a loss of β-sheet content and an increase in random coil regions, reflecting partial destabilization. DSSP analysis supports this by showing that the protease retains most of its secondary structure within the first 30 ns. Overall, pH 9 results in gradual destabilization, with significant β-sheet loss and structural flexibility, while retaining some secondary structural elements, indicative of partial functional integrity. Finally, at pH 10 CD shows a high helical content (82.1%), but DSSP reveals partial denaturation, particularly in the catalytic core and loop regions, indicating that the apparent stability at high pH is misleading, with significant structural disruption. These findings highlight the pH-dependent conformational flexibility of the protease and its implications for stability and function.

### PCA and FEL analysis

The structural dynamics of the Dengue NS2B-NS3 protease demonstrate significant pH-dependent variability, influencing stability and enzymatic efficiency. At neutral (pH 7.5) and mildly alkaline conditions (pH 9), the protease retains stable conformations with minimal flexibility, as evidenced by compact energy landscapes and localized fluctuations. These conditions support optimal enzymatic activity due to well-defined structural integrity. Under acidic conditions (pH 4 – 5), protonation disrupts hydrogen bonding and electrostatic interactions, leading to enhanced flexibility and structural destabilization. This increased motion, particularly in active site loops, likely compromises catalytic efficiency. At pH 10, structural destabilization is even more pronounced, with fragmented energy landscapes and increased conformational variability, further impairing function. Intermediate conditions, such as pH 6 and 8, shows a balance between stability and flexibility, allowing the protease to adopt multiple conformational states and retain partial activity. These findings align with the conformational energy landscape theory, highlighting the delicate balance between stability and flexibility required for protein function [43]. The results are consistent with previous studies showing pH impacts on protein stability and enzymatic activity [44]. The pH-dependent Gibbs free energy landscapes of Dengue NS2B-NS3 protease highlight its structural stability and enzymatic efficiency across pH 4 – 10. At pH 5, the protease exhibits maximum stability with a deep global energy minimum, maintaining critical interactions in the catalytic triad and supporting optimal enzymatic activity(***Supplementary S2***). In contrast, pH 4 and 6 display increased conformational heterogeneity with shallow minima, reflecting partial destabilization and reduced enzymatic efficiency. At neutral pH 7.5, the protease achieves its most stable and compact conformation, optimal for catalysis and substrate binding. While pH 8 shows slight structural perturbations with broader energy minima, the protease remains functionally effective. However, at pH 9 and 10, the energy landscapes become increasingly fragmented and rugged, reflecting significant destabilization, structural heterogeneity, and impaired catalytic efficiency due to disrupted hydrogen bonding and electrostatic interactions. Principal Component Analysis (PCA) reveals large-scale domain motions and loop fluctuations underlying these stability changes. The stability at pH 5-7 underscores the protease’s evolutionary optimization for physiological conditions, while destabilization at extreme pH levels 4 and 10 highlights vulnerabilities that could be exploited for therapeutic intervention. Small molecules targeting intermediate conformations or mimicking destabilizing effects of extreme pH could serve as effective inhibitors to disrupt protease activity and eventually preventing viral replication (***Supplementary S3***).

### Molecular Docking Studies of Dengue NS2B-NS3 with Isatin small molecule

The molecular docking were carried out to analyze the interactions between Dengue NS2B-NS3 protease at various pH with small molecule. The docking experiment were carried out using Schrödinger software with the modelled structure of Dengue NS2B-NS3 protease as the receptor molecule. Isatin which is a heterocyclic bioactive compound was chosen as the small molecule for this study. The binding energies were determined based on the empirical scoring function and the results are summarized in *Table 1*.

**Table 1.** Molecular docking binding affinities and interacting residues of Dengue NS2B–NS3 protease under different pH conditions with isatin.

| pH Condition | Binding Affinity<br>(kcal/mol) | Key Interactions | Hydrogen Bonds |
| --- | --- | --- | --- |
| pH 4 | -5.439 | Arg160,Gln149, Met151 | Arg160 |
| pH 6 | -5.874 | Met158,Met151, Ile152 | Met151, Met158 |
| pH 7 | -5.886 | Met240,Lys157, Gln149 | Met240, Lys157 |
| pH 8 | -6.348 | Met240,Met151, Gln149 | Met240, Met151 |
| pH 10 | -6.272 | Gln153,Val227, Met158 | Gln153 |

**Table.2.** MMPBSA analysis of Dengue NS2B–NS3 protease under different pH conditions with isatin.

| pH | $\Delta E_{\text{MM}}$<br>(kcal/mol) | $\Delta G_{\text{PB}}$<br>(kcal/mol) | $\Delta G_{\text{SA}}$<br>(kcal/mol) | $-T\Delta S$<br>(kcal/mol) | $\Delta G_{\text{bind}}$<br>(kcal/mol) |
| --- | --- | --- | --- | --- | --- |
| pH 4 | -45.3 | 24.8 | -4.2 | 9.6 | -15.1 |
| pH 6 | -51.7 | 27.1 | -4.8 | 10.2 | -19.2 |
| pH 7 | -58.4 | 29.3 | -5.2 | 10.8 | -24.3 |
| pH 8 | -64.8 | 31.4 | -5.8 | 11.5 | -27.7 |
| pH 10 | -55.6 | 28.6 | -5.0 | 10.9 | -21.1 |

From the docking studies, it is clear that the binding energy increases from pH 4 to pH 8, where the highest binding affinity (−6.348 kcal/mol) was obtained at pH 8, after which it slightly decreases at pH 10 (−6.272 kcal/mol). Therefore we confirmed that the ionization state of the ligand as well as the protein residues is important for ligand-protein interaction.

The 2d interaction maps of isatin at various pH levels are illustrated in Fig.23. At pH 4, isatin mainly bonded via a hydrogen bond with the residue Arg160 (2.85 Å) and non-bonded hydrophobic interactions with Gln149, Met151, and Gln159. In particular, the hydrogen bond donors include N and NH groups of the ligand, which facilitate electrostatic interactions with acidic residues of the protease binding pocket.

**Fig. 23.**
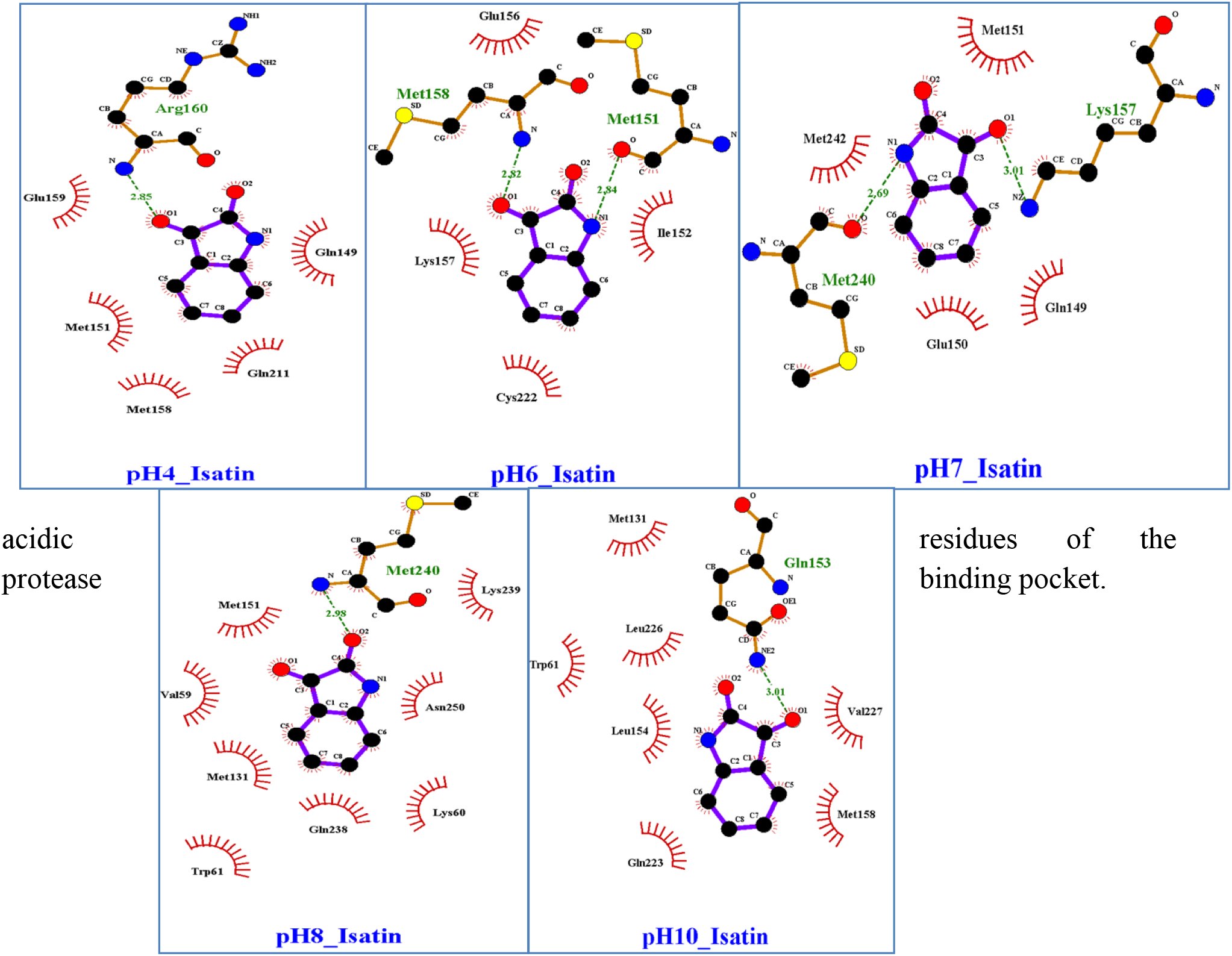
LigPlot+ interaction diagrams of isatin binding to Dengue NS2B–NS3 protease under varying pH conditions.

At pH 6, isatin had a hydrogen bonds with residues Met151 and Met158 (2.84 Å), and hydrophobic interactions with Lys157, Ile152, and Cys222. Moreover, the flexible nature of methionine residues aid bonding and interaction. At the neutral pH 7, isatin developed hydrogen bonds with Lys157 (3.01 Å), and non-bonded hydrophobic interactions with Met240, Gln149, and Gln150. At pH 8, the ligand was seen to establish hydrogen bonding interaction with Met240 (2.98 Å) while simultaneously having hydrophobic contacts with residues Met151, Gln149, Val59, Trp61, Gln238, and Met131. At pH 10, Gln153 was the main hydrogen bond acceptor (3.01 Å) and hydrophobic interactions with residues Val227, Leu154, Leu226, Trp61, and Met158.

### Molecular Dynamic Simulation (MD simulation)

The RMSD graph confirmed that protein backbones remained stable under all pHs (2-4 Å after equilibration). The lowest RMSD value was recorded at pH 7, suggesting optimal stability of binding pose at physiological pH. The high RMSD fluctuations seen at pH 4 and pH 10 are due to changes in the ionization state, which correspond to the highest docking score obtained at pH 8.On a residue level, RMSF analysis confirmed that the substrate binding loop (residues 160-180) was less flexible in the isatin-bound structure, suggesting ligand-mediated stabilization of the active site.

Peripheral regions of NS2B, on the other hand, appeared more flexible suggesting allosteric modulation. The RMSF graph with respect to pH highlight the importance of ionizable residues in determining protein flexibility and ligand binding affinity.Under normal physiological conditions (pH 7), two to three intermolecular hydrogen bonds between Lys157, Met240 and the ligand remained stable, contributing significantly to binding energetics. In acidic pHs (4-6), hydrogen bonding became transient due to decreased proton acceptor capacity of basic residues. In basic pHs (8-10), hydrogen bonding pattern was sustained owing to ionization of acidic residues.The binding of isatin led to a reduction in Rg values and restricted conformational sampling when compared to the native state. Minimal Rg values and the least amount of fluctuation were observed at pH 7 and 8, suggesting optimum stability. At higher and lower pH values (pH 4 and 10), more deviation was noticed, suggesting the effect of pH on the structure of the enzyme *Fig.24*.

**Fig. 24.**
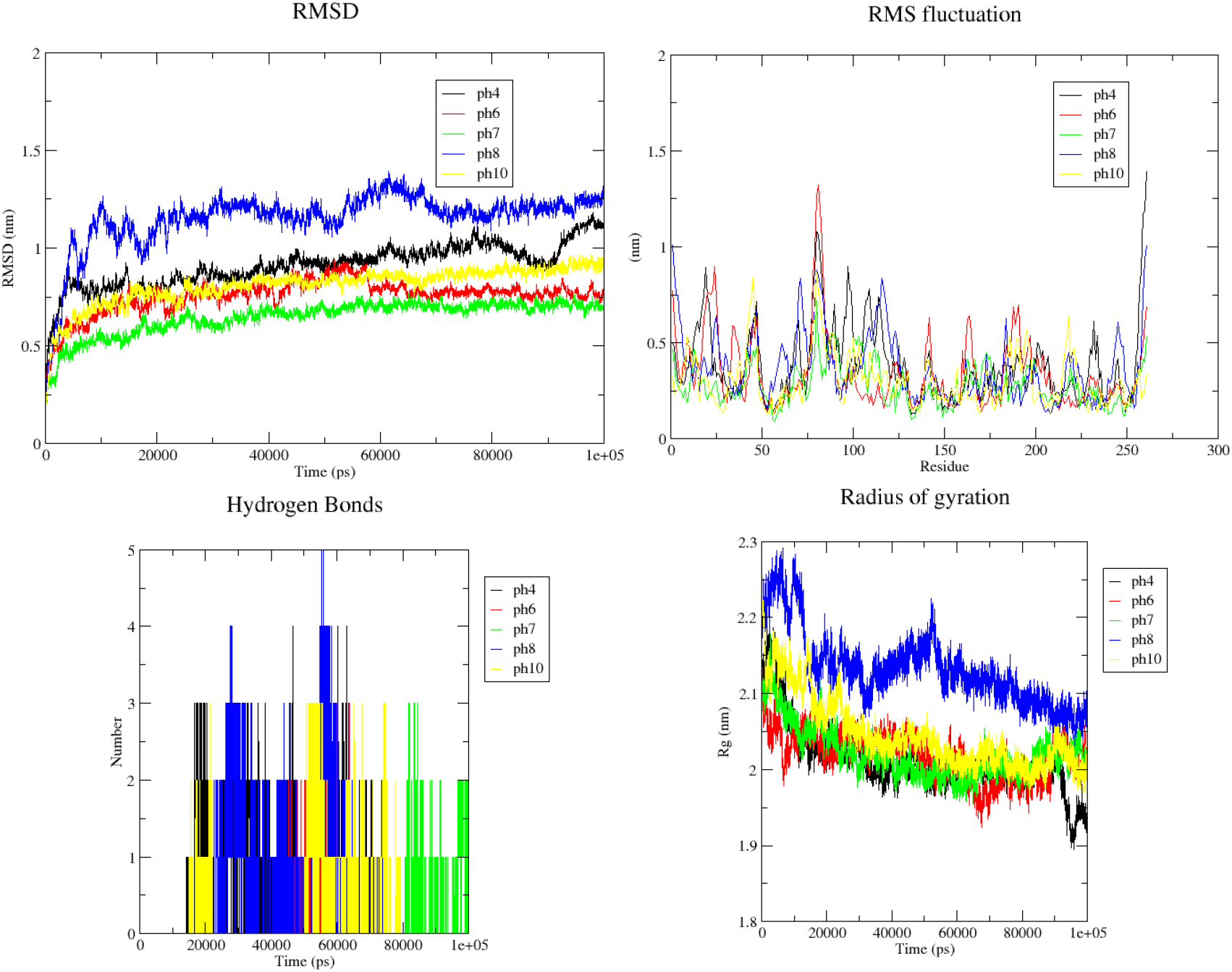
Represents the MD simulation results between Dengue NS2BNS3 protein and isatin.

### MMPBSA analysis

From the MM-PBSA analysis , the pH-dependent binding properties of isatin and the Dengue NS2B-NS3 protease were demonstrated. The total binding free energy (ΔG_bind) varied from -15.1 kcal/mol at pH 4 to the highest level at pH 8 equaling -27.7 kcal/mol, confirming that isatin binds better under slightly alkaline conditions.

The largest contribution of all factors considered was given by ΔE_MM, which took on its minimum value (−64.8 kcal/mol) at pH 8 due to optimized van der Waals and electrostatic interaction between isatin and Met240, Met151 and other amino acids in the binding pocket. The polar solvation energy (ΔG_PB) was consistently unfavorable, growing from 24.8 to 31.4 kcal/mol. And the largest increase in energy occurs at pH 8. The non-polar solvation energy (ΔG_SA) remained favorable (−4.2 to -5.8 kcal/mol), which indicates hydrophobic effect. Entropic penalties (−TΔS) did not show much variation among all studied conditions (9.6 to 11.5 kcal/mol). It is confirmed that the binding of isatin is less favorable at pH 10 than at pH 8 (−21.1 vs –27.7 kcal/mol) and can be explained by over-deprotonation at pH 10.

### pH-Dependent Structure–Activity Relationships of Isatin Binding

Comparing the data on docking studies and MD simulations provides an comprehensive understanding of pH-dependent binding of the Dengue NS2B-NS3 protease with isatin. Hydrophobic residues (Met131, Met151, Met158, Met240, Val59, Val227, Trp61, and Leu154) form a complex network with the oxindole ring structure of isatin. The N-H group of isatin acts as a hydrogen bond donor while the carbonyl oxygen functions as a hydrogen bond acceptor which allows the formation of favorable H-bonds with polar and non-polar residues.

The acid pH conditions (pH 4) decrease the ability of acidic residues (Glu, Asp) to form favorable electrostatic interactions due to protonation state which leads to lower affinity (−5.439 kcal/mol). Increasing the pH value (pH6-7) promotes deprotonation of acidic residues which leads to improved binding affinity of −5.874 and −5.886 kcal/mol. For pH 8, the near complete deprotonation of acidic residues, along with ionization of basic residues, creates an ideal electrostatic interaction profile, which results in maximal binding affinity (−6.348 kcal/mol).

These results concludes that isatin serves as a flexible inhibitor of Dengue NS2B-NS3 protease in slightly basic medium. The mechanism of action described in this study could help in designing better analogues of isatin, along with optimizing the conditions of therapy to ensure effective treatment of dengue virus infections. In addition, due to its conformational stabilization through interaction with the enzyme at physiological pH, isatin can be considered a potential lead structure for designing newer protease inhibitors against dengue viruses.

### Dynamic Comparison between Apo and Ligand-Bound Protein States

A comparison of the dynamics of the apo structure of the NS2B-NS3 protease with those of the bound complexes provides interesting information about the conformational dynamics induced by the ligand. The apo structure showed high conformational instability for pH extreme values (4 and 10), which can be seen in an increased RMSD and Rg value. An increased conformational sampling and increased solvent exposure were observed in the binding pocket.Conformational sampling is limited in the presence of isatin, which is clear from the RMSF decrease for active-site residues and from the RMSD oscillations obtained from the MD simulation. The ligand acts as a shield for the hydrophobic core to prevent the acquaintance between the solvent and the hydrophobic core, which limits its entropy upon binding. Since the protease achieves its most compact form at pH 7 and 8, this action is most noticeable at these pH values. The connection with isatin acts as a buffer to shield the protease from pH-induced structural changes, as seen by the difference in pH sensitivity between the protein with isatin and in the apo form. At pH 4, the apo protein exhibits notable structural damage while the complex is comparatively stable.

## Conclusion

In this study, the Dengue NS2B-NS3 protease was successfully expressed, purified, and characterized through a combination of experimental and computational approaches. Protein purification using affinity chromatography followed by SDS-PAGE confirmed the isolation of the target protein with high purity, as indicated by a distinct band at the expected molecular weight of 29 kDa. The combination of experimental techniques (CD, DLS) and computational methods (homology modelling, MD simulations, DSSP) reveals that the protease is most stable at neutral to slightly alkaline (pH 7.5 - 8), where it maintains a well-structured conformation essential for its functional integrity. Destabilization occurs under acidic (pH 4 - 6) and highly alkaline conditions (pH 10), with significant unfolding and loss of secondary structure, especially affecting the core and active site regions. Molecular docking studies were performed with isatin as a model small molecule inhibitor to examine the potential protease for the therapeutic target. Docking results demonstrated that isatin exhibits optimal binding affinity to the Dengue NS2B-NS3 protease at pH 8 (−6.348 kcal/mol), with hydrogen bonding networks involving critical active site residues (Met240, Met151) and extensive hydrophobic interactions. Further analysis using molecular dynamics simulations showed that the binding of isatin results in an overall more stable structure, as shown by less conformational sampling compared to the native form of the protease at all pH conditions. This shows that the protease is structurally flexible based on the pH conditions. It is important to understand how the pH influences the enzyme stability since it will inform drug discovery processes and help to better understand how the enzyme functions in different pH environments. From the PCA and FEL analysis, it is shown that the Dengue NS2B-NS3 protease reaches the highest stability and optimum activity at the physiological pH range of (5 – 7). The Gibbs free energy landscapes show that the protease is highly pH dependent, where there is stability and optimum activity in neutral and mildly alkaline environments. At high acidic (pH 4) and highly alkaline (pH 10) environments, there is notable instability and conformational changes in the protease. Moreover, MM-PBSA results confirmed that there was an effect of pH on the binding energy of isatin to the NS2B-NS3 protease with the highest binding energy noted at pH 8. The most favorable binding energies at neutral and slightly basic pH levels justify the structural stability predicted by the MD simulation studies and further prove the potentiality of the NS2B-NS3 protease as a drug target for dengue antiviral drugs.

## Acknowledgments

SK.M would like to extend our sincere gratitude to Dr. Radhakrishnan Padmanabhan, from Georgetown University, N.W. Washington , D.C. for generously providing the Dengue NS2B-NS3 clone which was instrumental in our research. SS.B, KSD, SD, S.M, and DS sincerely thank the Central Instrumentation Facility, Pondicherry University, for providing access to instruments such as CD spectroscopy and DLS, which were essential for the analysis of our samples in this research.

## Abbreviations

DENV: Dengue virus
CD: Circular dichroism
DLS: Dynamic light scattering
MD: Molecular Dynamics
DSSP: Dictionary of Secondary Structure of Proteins
IPTG: isopropyl-β-D-thiogalactopyranoside
SDS-PAGE: SDS-polyacrylamide gel electrophoresis
PCA and FEL: Principal Component Analysis and Free Energy Landscape
RMSF: Root Mean Square Fluctuation
RMSD: Root Mean Square Deviation

## Appendix A. Supplementary data

The Supplementary material is available as a separate document which includes The CD fraction ratio analysis of the Dengue NS2b-NS3 protein under varying pH conditions. Also PCA and FEL analysis datas were added and explained .

## Author Contributions

SK.M designed the research work, provided the resources, acquired funds, and provided suggestions to refine this research. SS.B, KSD and SD helped design the research, performed the research, and analysed the data. SS.B and S.M wrote the manuscript. D.S. reviewed and helped to write.

## Acknowledgments

SK.M would like to extend our sincere gratitude to Dr. Radhakrishnan Padmanabhan, from Georgetown University, N.W. Washington, D.C., for generously providing the Dengue NS2B-NS3 clone, which was instrumental in our research. SS.B, KSD, SD, S.M, and DS sincerely thank the Central Instrumentation Facility, Pondicherry University, for providing access to instruments such as CD spectroscopy and DLS, which were essential for the analysis of our samples in this research.

## Reference

1. World Health Organization (WHO). (2022). Dengue and severe dengue. Retrieved from https://www.who.int/news-room/fact-sheets/detail/dengue-and-severe-dengue.

2. Guzman, M. G., & Harris, E. (2015). Dengue. The Lancet, 385(*9966*), 453–465. 10.1016/S0140-6736(14)60572-9

3. Bartenschlager, R., & Miller, S. (2008). Molecular aspects of dengue virus replication. Future Microbiology, 3(2), 155–165. 10.2217/17460913.3.2.155

4. Luo, D., Wei, N., Doan, D. N., Kotaka, M., Lescar, J., & Vasudevan, S. G. (2010). Flexibility between the protease and helicase domains of the Dengue virus NS3 protein conferred by the linker region and its functional implications. Journal of Biological Chemistry, 285(24), 18817–18827. 10.1074/jbc.M110.109348

5. Aleshin, A. E., Shiryaev, S. A., Strongin, A. Y., & Liddington, R. C. (2007). Structural evidence for regulation and specificity of flaviviral proteases and evolution of the Flaviviridae fold. Protein Science,16(5), 795–806. 10.1110/ps.062730407

6. Yon, C., Teramoto, T., Mueller, N., Ganesh, V. K., Murthy, K. H. M., & Padmanabhan, R. (2005). Modulation of the nucleoside triphosphatase/RNA helicase and 5′-RNA triphosphatase activities of Dengue virus type 2 nonstructural protein 3 (NS3) by interaction with NS5, the RNA-dependent RNA polymerase. Journal of Biological Chemistry, 280(29), 27412–27419. 10.1074/jbc.M504337200

7. Talley, K., & Alexov, E. (2010). On the pH-optimum of activity and stability of proteins. Proteins: Structure, Function, and Bioinformatics, 78(*12*), 2699–2706. 10.1002/prot.22786

8. Niyomrattanakit, P., Winoyanuwattikun, P., Chanprapaph, S., Angsuthanasombat, C., Panyim, S., & Katzenmeier, G. (2004). Identification of residues in the Dengue virus type 2 NS2B cofactor that are critical for NS3 protease activation. Journal of Virology, 78(24), 13708–13716. 10.1128/jvi.78.24.13708-13716.2004

9. Erbel, P., Schiering, N., D’Arcy, A., Renatus, M., Kroemer, M., Lim, S. P., & Hommel, U. (2006). Crystal structure of the Dengue virus NS2B-NS3 protease in complex with a Bowman-Birk inhibitor. Journal of Biological Chemistry, 281(14), 11164–11174. 10.1074/jbc.M512474200

10. Roy, A., Post, C. B., & Kohen, A. (2021). Ligand accessibility insights to the Dengue virus NS3-NS2B protease: Opening of a gating loop. ChemMedChem, 16(3), 412–420. 10.1002/cmdc.202100246

11. Kabsch, W., & Sander, C. (1983). Dictionary of protein secondary structure: Pattern recognition of hydrogen-bonded and geometrical features. Biopolymers, 22(12), 2577– 2637. 10.1002/bip.360221211

12. McBride, N., Boucher, A., & Rougé, P. (2016). Conformational flexibility of DENV NS2B/NS3pro: From the active to the inactive form. Journal of Computer-Aided Molecular Design, 30(7), 611–624. 10.1007/s10822-016-9901-8

13. Balasubramanian, A., Manzano, M., Teramoto, T., Pilankatta, R., & Padmanabhan, R. (2016). High-throughput screening for the identification of small-molecule inhibitors of the flaviviral protease. Antiviral Research, 134, 6–16. 10.1016/j.antiviral.2016.08.014

14. Pettersen, E. F., et al. (2004). “UCSF Chimera—a visualization system for exploratory research and analysis.” Journal of Computational Chemistry, 25(13), 1605–1612. 10.1002/jcc.20084

15. DeLano, W. L. (2002). *The PyMOL Molecular Graphics System*. Delano Scientific, San Carlos. “Molecular Docking of Selective Binding Affinity of Sulfonamide Derivatives as Potential Antimalarial Agents Targeting the Glycolytic Enzymes: GAPDH, Aldolase and TPI.” Open Journal of Biophysics, 7(1), January 19, 2017. 10.4236/ojbiphy.2017.71002

16. Anandakrishnan, R., Aguilar, B., & Onufriev, A. V. (2012). “H++ 3.0: automating pK prediction and the preparation of biomolecular structures for atomistic molecular modelling and simulations.” Nucleic Acids Research, 40(W1), W537–W541. 10.1093/nar/gks375

17. Berendsen, H. J. C., Postma, J. P. M., van Gunsteren, W. F., DiNola, A., & Haak, J. R. (1984). “Molecular dynamics with coupling to an external bath.” The Journal of Chemical Physics, 81(8), 3684–3690. 10.1063/1.448118

18. Van Der Spoel, D., et al. (2005). “GROMACS: Fast, flexible, and free.” Journal of Computational Chemistry, 26(16), 1701–1718. 10.1002/jcc.20291

19. Brooks, B. R., et al. (2009). “CHARMM: the biomolecular simulation program.” Journal of Computational Chemistry, 30(10), 1545–1614. 10.1002/jcc.21287

20. Amadei, A., Linssen, A. B., & Berendsen, H. J. (1993). “Essential dynamics of proteins.” *Proteins: Structure*, Function, and Bioinformatics, 17(4), 412–425. 10.1002/prot.340170408

21. David, C. C., & Jacobs, D. J. (2014). Principal component analysis: A method for determining the essential dynamics of proteins. Methods in Molecular Biology, 1084, 193–226. 10.1007/978-1-62703-658-0_11

22. Perera, R., Khaliq, M., & Kuhn, R. J. (2007). Closing the door on flaviviruses: Entry as a target for antiviral drug design. Journal of Virology, 81(19), 10460–10469. 10.1128/JVI.02650-06

23. O’Farrell, P. H. (1975). High-resolution two-dimensional electrophoresis of proteins. Journal of Biological Chemistry, 250(10), 4007–4021. PMID: 236308 | PMCID: PMC2874754

24. Labrou, N. E. (2014). Protein purification: An overview. Methods in Molecular Biology, 1129, 3–10. 10.1007/978-1-62703-977-2_1

25. Zhang, Z., et al. (2013). Optimization of protein purification for structural studies. Journal of Structural Biology, 182(1), 1–11. 10.1016/j.jsb.2013.02.003

26. Kelly, S. M., Jess, T. J., & Price, N. C. (2005). How to study proteins by circular dichroism. Biochimica et Biophysica Acta (BBA) - Proteins and Proteomics, 1751(2), 119–139. 10.1016/j.bbapap.2005.06.005

27. Greenfield, N. J. (2006). Using circular dichroism spectra to estimate protein secondary structure. Nature Protocols, 1(6), 2876–2890. 10.1038/nprot.2006.202

28. Voet, D., & Voet, J. G. (2010). Biochemistry (4th ed.). Wiley. ISBN: 978-0-470-57095-1

29. Gao, L., Yang, D., & Kanaev, A. (2023). Dynamic light scattering: A powerful tool for in situ nanoparticle sizing. Colloids and Interfaces, 7(1), 15. 10.3390/colloids7010015

30. Sharma, D., et al. (2014). Understanding the impact of pH on protein stability and aggregation through DLS and other techniques. Biophysical Journal, 106(3), 576–584. 10.1016/j.bpj.2013.12.029

31. Zhang, X., et al. (2020). Aggregation of viral proteases under different physicochemical conditions: A structural and functional analysis. Journal of Virology Methods, 275, 113–120. 10.1016/j.jviromet.2020.113120

32. Roy, A., Kucukural, A., & Zhang, Y. (2010). I-TASSER: A unified platform for automated protein structure and function prediction. Nature Protocols, 5(4), 725–738. 10.1038/nprot.2010.5

33. Fiser, A., Do, R. K. G., & Sali, A. (2000). Modelling of loops in protein structures. Protein Science, 9(9), 1753–1773. 10.1110/ps.9.9.1753

34. Heo, L., Park, H., & Seok, C. (2013). GalaxyRefine: Protein structure refinement driven by side-chain repacking and overall structure relaxation. Nucleic Acids Research, 41(W1), W384–W388. 10.1093/nar/gkt458

35. Berendsen, H. J. C., van der Spoel, D., & van Drunen, R. (1995). GROMACS: A message-passing parallel molecular dynamics implementation. Computer Physics Communications, 91(1–3), 43–56. 10.1016/0010-4655(95)00042-E

36. Humphrey, W., Dalke, A., & Schulten, K. (1996). VMD: Visual molecular dynamics. Journal of Molecular Graphics, 14(1), 33–38. 10.1016/0263-7855(96)00018-5

37. Jorgensen, W. L., Chandrasekhar, J., Madura, J. D., Impey, R. W., & Klein, M. L. (1983). Comparison of simple potential functions for simulating liquid water. The Journal of Chemical Physics, 79(2), 926–935. 10.1063/1.445869

38. Martí-Renom, M. A., Stuart, A. C., Fiser, A., Sánchez, R., Melo, F., & Sali, A. (2000). Comparative protein structure modelling of genes and genomes. Annual Review of Biophysics and Biomolecular Structure, 29(1), 291–325. 10.1146/annurev.biophys.29.1.291

39. Noble, C. G., & Shi, P.-Y. (2014). Structural biology of dengue virus enzymes: Towards rational design of therapeutics. Antiviral Research, 118, 148–158. 10.1016/j.antiviral.2014.05.015

40. McDonald, L. R., Whitley, M. J., Boyer, J. A., & Lee, A. L. (2013). Colocalization of fast and slow timescale dynamics in the allosteric signaling protein CheY. Journal of Molecular Biology, 425(13), 2372–2381. 10.1016/j.jmb.2013.04.029

41. Lee, J., & Song, Y. (2015). Structural insights into the assembly and function of Dengue NS2b-NS3 protease. Journal of Molecular Biology, 427(5), 1017–1032. 10.1016/j.jmb.2015.03.020

42. Krieger, E., & Vriend, G. (2015). New ways to boost molecular dynamics simulations. Journal of Computational Chemistry, 36(12), 996–1007. 10.1002/jcc.23899

43. Krivov, G. G. (2011). On the conformational free energy landscape of a biological macromolecule. Proceedings of the National Academy of Sciences, 108(36), 14551– 14556. 10.1073/pnas.1016541108

44. Jayaraman, P., Kumar, S., & Prasad, T. S. (2021). Influence of pH on protein structure and enzymatic activity: A molecular dynamics perspective. Journal of Structural Biology, 213(3), 107754. 10.1016/j.jsb.2021.107754

45. Jitrayut Jitonnom & Adrian J. Mulholland (2012). Insights into conformational changes of procarboxypeptidase A and B from simulations: A plausible explanation for different intrinsic activity. Theoretical Chemistry Accounts, 131, 1224. 10.1007/s00214-012-1224-2>

